# Effects of knockdown of autophagy pathway genes on *C. elegans* longevity are highly condition dependent

**DOI:** 10.1101/2025.07.28.667102

**Authors:** Kuei Ching Hsiung, Hannah Chapman, Xiaoya Wei, Xiaoyu Sun, Isadora Rawlinson, David Gems

**Affiliations:** Institute of Healthy Ageing, and Research Department of Genetics, Evolution and Environment, University College London, London, UK

**Keywords:** aging, autophagy, condition dependence, germline, insulin/IGF-1 signaling, nematode

## Abstract

Autophagy is proposed to protect against aging by clearing damaged cellular constituents. In line with this several life-extending interventions in model organisms show some degree of autophagy dependence. In *C. elegans*, inhibiting autophagy can shorten, lengthen or have no effect on lifespan. Differences between published findings likely reflect variability in experimental conditions. Here we investigate the condition dependence of effects on lifespan of RNA-mediated interference (RNAi) knockdown of autophagy pathway components. Effects on interventions causing a strong Age (increased lifespan) phenotype were examined, mainly mutation of *daf-2* (insulin/IGF-1 receptor). Factors varied included *daf-2* mutant allele class, *atg* gene and temperature. Effects on lifespan of *atg* RNAi proved to be highly condition dependent. Notably, for most *atg* genes tested lifespan was not usually reduced more in the long-lived mutant than in the wild-type control. Greater suppression was seen at 20°C for certain *atg* genes with *daf-2(e1368)* but not *daf-2(e1370)*. At 25°C, little reduction in lifespan was seen. However, *atg-18* knockdown behaved differently, suppressing *daf-2* Age under all conditions, suggesting possible pleiotropic action. Condition dependent effects were also seen with respect to *glp-1* (reduced germline development) Age. Furthermore, presence of 5-fluoro-2’-deoxyuridine (FUDR) at a high concentration caused knockdown of several *atg* genes to increase lifespan in wild-type *C. elegans*. Thus, depending on experimental conditions, *atg* knockdown can increase, decrease or have no effect on *daf-2* Age. The lack of suppression of *daf-2* Age by *atg* RNAi under most conditions questions the importance of autophagy for this phenotype. Moreover, condition dependence of effects creates a risk of possible condition selection bias.

## Introduction

A long-standing theory is that senescence (aging) is largely a consequence of the accumulation of random molecular damage caused by, among other things, reactive oxygen species (Beckman and Ames, 1998; Harman, 1956; Murphy, 2023; Shore and Ruvkun, 2013; Szilard, 1959). This view predicts that mechanisms of somatic maintenance, particularly those that prevent accumulation of damaged cellular constituents, counteract the aging process. One somatic maintenance function viewed as a potential longevity-assurance process is autophagy (specifically macroautophagy), which effects lysosome-dependent degradation of cellular constituents, including damaged matter (Aman et al., 2021).

The possible anti-aging role of autophagy has been extensively tested in the short-lived nematode *Caenorhabditis elegans*, and supporting evidence found with respect to several life-extending interventions, including reduced insulin/IGF-1 signaling (IIS), reduced germline signaling, and dietary restriction (Hansen et al., 2018). These studies were principally performed in the 2000s; however, during the same period, falsification tests of the molecular damage theory, particularly that relating to oxidative damage, led to growing uncertainty about its validity (Gems and Doonan, 2009; Perez et al., 2009; Shields et al., 2021). Meanwhile, an alternative theoretical framework emerged, based on the evolutionary theory of aging (Arnold and Rose, 2023; Williams, 1957), arguing that senescence is largely the consequence of genetically-determined, programmatic mechanisms (Blagosklonny, 2006; de Magalhães and Church, 2005; Gems, 2022; Maklakov and Chapman, 2019), and very much so in *C. elegan*s (Gems and de la Guardia, 2013; Gems et al., 2021; Pires da Silva et al., 2024).

One form of programmatic aging involves costly programs: genetically-determined processes that consume or degrade somatic tissues as a by-product of wider, fitness-promoting processes (Gems and Kern, 2024; Gems et al., 2021). In some cases this involves biomass repurposing, where biomass of one tissue is broken down by autophagic processes to release molecular constituents to support functions in another. This occurs to a high degree in semelparous organisms in the process of reproductive death (suicidal reproductive effort), as in many monocarpic plants, and semelparous fish such as Pacific salmon (Gems et al., 2021).

Several lines of evidence support the hypothesis that reproductive death occurs in *C. elegans* hermaphrodites (Gems et al., 2021; Kern et al., 2023). This includes a putative costly program in which intestinal biomass is broken down and repurposed to support production of yolk, that is then vented to support larval growth, leading to intestinal atrophy (a senescent pathology) (Ezcurra et al., 2018; Kern et al., 2021; Sornda et al., 2019). Notably, such biomass repurposing is supported by autophagy, as evidenced by deceleration of intestinal atrophy and yolk pool formation when autophagy is inhibited (Benedetto and Gems, 2019; Ezcurra et al., 2018). Thus, in this particular context autophagy appears to be promoting rather than preventing senescence. It is by now clear that autophagy can enhance as well as inhibit the development of pathologies in *C. elegans*, including senescent ones (Kang et al., 2007).

These developments warrant a careful reassessment of the evidence that autophagy is protective against aging in *C. elegans*. In this study we reexamine the question of whether the longevity of *daf-2* insulin/IGF-1 receptor mutants is autophagy dependent. Here the principal form of past evidence involves epistasis tests, where effects on lifespan of reduction of function of genes encoding proteins effecting autophagy is compared in the wild type (N2) and *daf-2* mutants. Findings from 9 previous studies involving 46 epistasis experiments are summarised in Table S1. Although it is widely believed that autophagy is essential for *daf-2* mutant longevity (Meléndez et al., 2003), scrutiny of the results of past tests raises doubts about the strength of this claim.

Careful consideration of these prior studies identifies six distinct limitations, as follows. (1) A life-shortening effect of inhibition of autophagy does not necessarily indicate its role in the normal aging process, or in *daf-2* mutant longevity. (2) If autophagy is inhibited during development as well as adulthood, a life-shortening effect could result from disruption of normal development. (3) The claim that *daf-2* longevity is autophagy dependent requires evidence that the life-shortening effect in *daf-2* is significantly greater than in wild type, e.g. by using Cox proportional hazard analysis, and this is rarely performed. (4) If effects of reducing autophagy are condition dependent, this introduces a potential bias: a risk that investigators might unwittingly select conditions where the data generated supports a role of autophagy in longevity - what may be referred to as *condition selection bias*. Potential determinative conditions that have varied across studies include choice of autophagy-determining gene to inhibit, of *daf-2* mutant allele, and ambient temperature. (5) 5-fluoro-2’-deoxyuridine (FUDR) is sometimes used to facilitate *C. elegans* lifespan assays by preventing progeny hatching; this could potentially affect test outcomes, as shown in other contexts (Aitlhadj and Sturzenbaum, 2010; Anderson et al., 2016; Burnaevskiy et al., 2018; Van Raamsdonk and Hekimi, 2011; Zhao et al., 2019). (6) A final, straightforward issue is whether a given finding proves to be reproducible in subsequent reports under, seemingly, the same conditions.

Of 46 prior tests (Table S1) only 4 present clear evidence that reducing autophagy shortens lifespan more in *daf-2* than in the wild type. Regarding one of these four instances, the effect of *bec-1* RNAi on *daf-2(e1370)* at 15°C (Meléndez et al., 2003), a subsequent study did not replicate this finding (Hars et al., 2007). In two other cases the weaker *daf-2(mu150)* allele was used (Hansen et al., 2008; Patel et al., 2008). In two further studies where large reductions in *daf-2* lifespan were observed (Chang et al., 2017; Minnerly et al., 2017) knockdown of *atg-18* was used. In a 2009 study where effects of knockdown of diverse autophagy genes was tested, *atg-18* was one of only 2/14 genes where adult-limited RNAi significantly reduced lifespan in the wild type (Hashimoto et al., 2009), suggesting possible *atg-18* idiosyncrasy. The 2009 study includes more than half of all of the prior tests (28/46); strikingly, in only 1/14 genes (*atg-4.1*) did adult-limited RNAi significantly reduce *daf-2(e1370)* lifespan, while for 3/14 genes (*atg-9*, *bec-1* and *unc-51*) it *increased daf-2* lifespan (Hashimoto et al., 2009). One possible reason for the lack of observed life-shortening effects given autophagy knockdown is its use of FUDR (high concentration), unlike other studies.

Here we assess the condition-dependence of the effects of RNAi knockdown of autophagy genes on *C. elegans* longevity. To this end we have tested the effects of RNAi of six genes in the autophagy pathway on longevity in two different *daf-2* mutants, and a *glp-1* germlineless mutant, at two temperatures. We have also assessed effects of FUDR and prevention of bacterial infection. Our results provide a robust foundation of evidence relating to possible autophagy dependence of *daf-2* and *glp-1* longevity. They suggest a weak and highly-condition dependent contribution of autophagy to *daf-2* and *glp-1* Age. This suggests that prevention of damage accumulation by autophagy is, at most, a minor determinant of *daf-2* longevity.

## Results

### Effects of *atg* RNAi on *daf-2* Age vary with *daf-2* allele

We first tested whether effects on lifespan of inhibiting autophagy depend upon *daf-2* allele severity. Two *daf-2* mutants were examined. *daf-2(e1368)* is a class 1 (less pleiotropic) mutant where adults show normal behavior at both 20°C and 25°C, while *daf-2(e1370)* is a class 2 (more pleiotropic) mutant where at 25°C adults become paralyzed and cease feeding (Gems et al., 1998). Knockdown of autophagy was performed by RNAi from the L4 (late larval) stage, to exclude possible confounding, life-shortening disruption of normal development. RNAi was performed for six genes at several stages of the autophagy process: initiation (*atg-13*), membrane nucleation (*atg-9*, *bec-1*), phagophore formation (*atg-2*, *atg-18*) and elongation (*atg-4.1*) (Figure 1A). 5/6 genes were those examined in our previous study (Ezcurra et al., 2018), with *bec-1* added because it was the subject of several earlier studies (Table S1).

**Figure 1.**
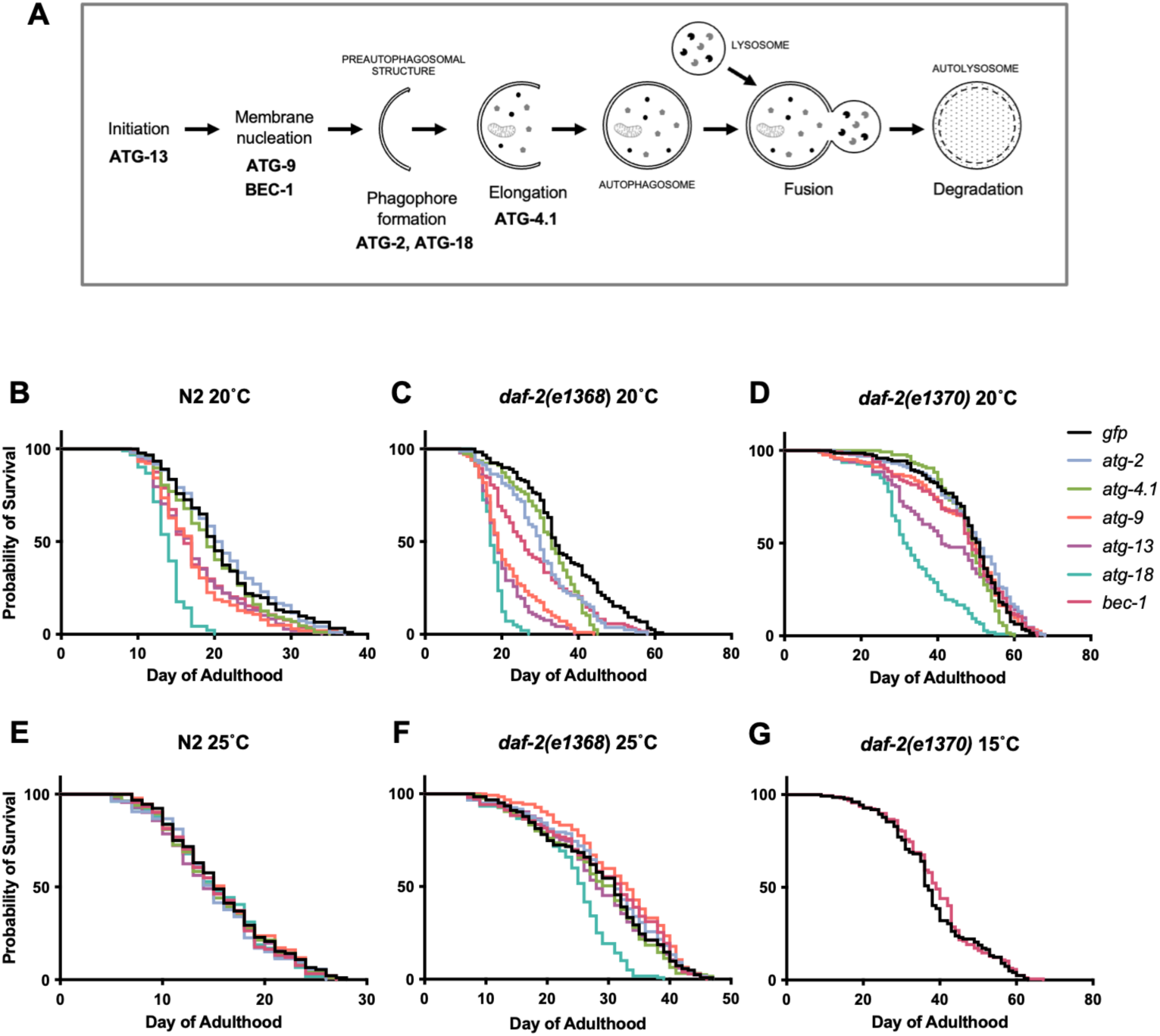
Effects of *atg* RNAi on longevity are highly variable and condition dependent. (A) The autophagy pathway, and genes tested in this study. (B-D) Effects at 20°C. (B) Effects on N2 (wild type). (C) Effects on *daf-2(e1368)*. (D) Effects on *daf-2(e1370)*. (E, F) Effects at 25°C. (E) Effects on N2. (F) Effects on *daf-2(e1368)*. (G) Effects of *bec-1* RNAi on *daf-2(e1370)* at 15°C. (B-G summed data, *N* = 2; for individual trial data and statistical comparisons, see Table S2, S5.

A methodological note: for tests of effects of a given intervention on *C. elegans* lifespan an often-applied standard is to include 3 biological replicates. This is true of several recent studies where the effect of knockdown of a single *atg* gene on *daf-2* longevity was studied (Minnerly et al., 2017; Wilhelm et al., 2017; Yang et al., 2024). However, given that the present condition dependence study effectively performs this test in 18 different ways, involving RNAi of 6 *atg* genes, 2 *daf-2* mutants and 2 temperatures, *N* = 2 biological replicates were judged to be sufficient to draw robust conclusions; similarly, an earlier study of RNAi 14 *atg* genes under two conditions used 2-3 biological replicates (Hashimoto et al., 2009); for an overview of the number of replicate trials performed in previous studies, see Table S1.

Trials were performed at 20°C, and used *gfp* RNAi as a negative control (with induction of an RNAi response). In wild-type *C. elegans* (N2), statistically significant reductions were sometimes seen: *atg-9, atg-13* and, particularly, *atg-18* RNAi shortened lifespan in both trials (summed data, *N* = 2, means:-13.7%, *p* < 0.0001,-12.4%, *p* = 0.0002,-31%, *p* < 0.0001, respectively; log rank test) (Figure 1B, Table S2; for all raw mortality data, see Supplementary Dataset 1). For survival curves comparing RNAi effects of individual *atg* genes on the three genotypes, see Figure 2.

**Figure 2.**
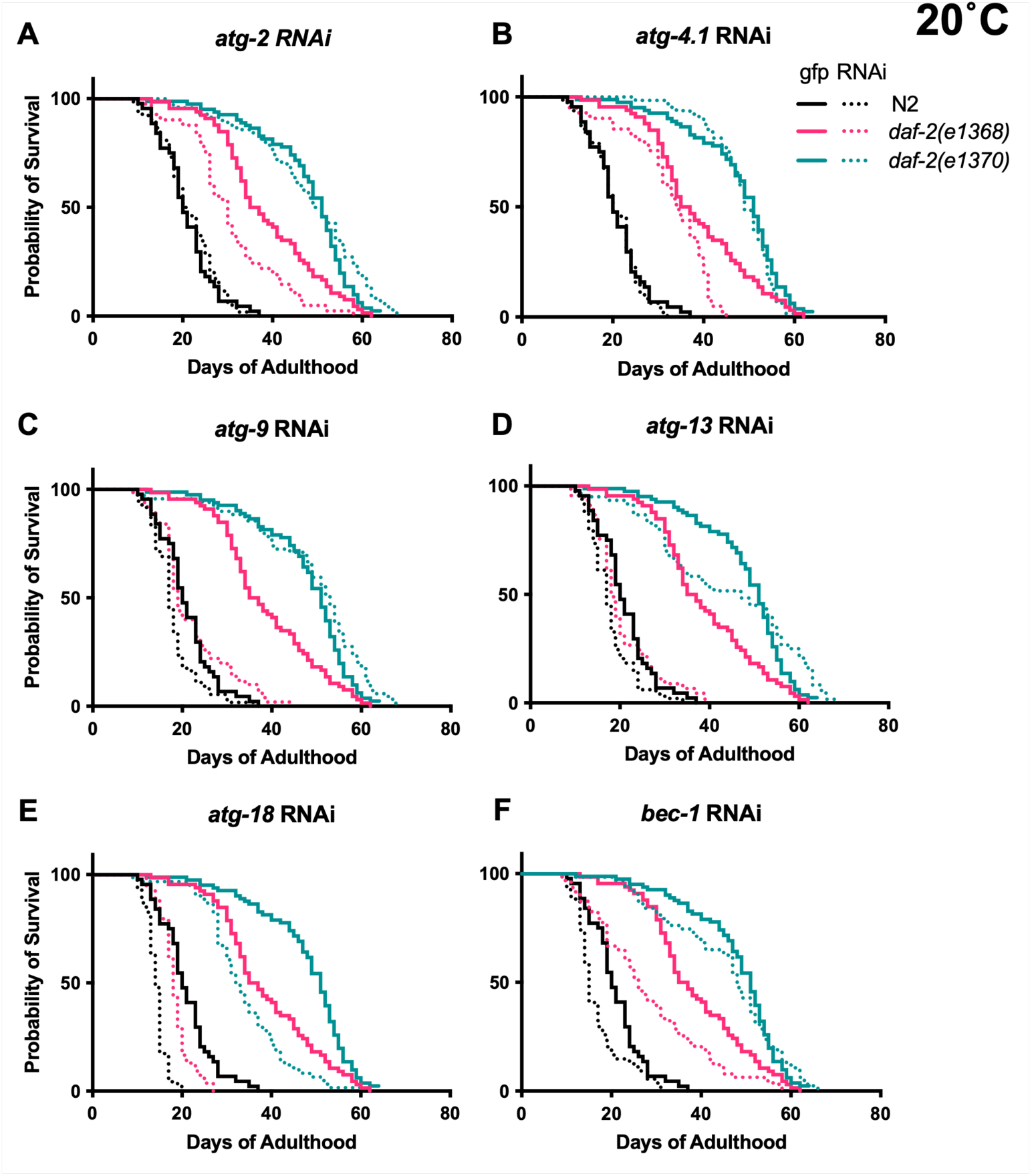
Degree of suppression of *daf-2* longevity by *atg* gene RNAi differs greatly between *daf-2* alleles and *atg* RNAi treatments (20°C), cf. Figure 1B-D (same data). Summed data, *N* = 2; for individual trials and statistical comparisons, see Table S2. All trials were performed in parallel, i.e. all lifespans (summed or in each trial) are directly comparable.

RNAi of all six *atg* genes consistently reduced lifespan in the *daf-2(e1368)* mutant, with effects ranging from-10.5% (*p* = 0.0009) for *atg-4.1* to-50.4% (*p* < 0.0001) for *atg-18* (Figure 1C, Table S2). To assess whether *atg* RNAi reduced lifespan more in *daf-2* mutants than in the wild type, Cox proportional hazard (CPH) analysis was used. This showed a significantly greater effect in *daf-2(e1368)* for 4/6 genes (exceptions: *atg-4.1*, *bec-1*) (Table S2). These findings are broadly consistent with the earlier observation that *bec-1* and *vps-34* RNAi shortened the lifespan of the *daf-2(mu150)* class 1 mutant but not of N2 at 20°C (CPH analysis not performed) (Hansen et al., 2008) (though in the present study *bec-1* RNAi did also reduce N2 lifespan, Table S2).

To test whether *atg* RNAi is able to fully suppress *daf-2(e1368)* Age, the lifespans of N2 and *daf-2* subjected to *atg* RNAi were compared. Under all six *atg* RNAi conditions, *daf-2(e1368)* still significantly increased mean lifespan, from +15.7% (*p* = 0.0031) for *atg-13* to +60.9% (*p* < 0.0001) for *atg-4.1* (summed data, 20°C, Table S2). This could imply either that *daf-2(e1368)* Age is not fully autophagy dependent, or that it is but autophagy is not fully suppressed by the RNAi treatments used.

In *daf-2(e1370)*, *atg* RNAi had far more modest effects on lifespan than in *daf-2(e1368)*. Summed data showed significant reductions in lifespan resulting from RNAi of *atg-9*, *atg-13* and *atg-18* only, with the latter again causing the largest reduction:-8.3% (*p* = 0.012),-17.8% (*p* = 0.0034), and-29.2% (*p* < 0.0001), respectively (Figure 1D, Table S2). However, effects were in no instance significantly greater than in N2 (CPH analysis, Table S2), thus failing to provide evidence that *e1370* longevity is mediated by autophagy. Taken together, these results suggest that at 20°C autophagy contributes to longevity in weaker but not stronger *daf-2* mutants, possibly in class 1 but not class 2 mutants.

RNAi effects varied between *atg* genes, with *atg-2* and *atg-4.1* RNAi effects weaker, and *atg-18* RNAi effects generally stronger than the rest. One possible cause of this variability is differences between the plasmid vectors used for RNAi in terms of how efficiently they destroy their target mRNA. To investigate this possibility, RNAi of each *atg* gene was performed in N2 animals, and transcript levels measured by RT–qPCR and analyzed using the ΔΔCt method, normalized to a *gfp* RNAi control, with fold changes calculated as 2^−ΔΔCt (*N* = 4 independent trials). Effects of RNAi on mRNA level varied greatly, from no detected reduction in *atg-2* to an 86% reduction in *atg-18* (Fig. S1A; Table S3; Supplementary dataset 2). Pairwise comparisons of ΔΔCt values showed that, *atg-2* aside, there were no significant differences between RNAi treatment effects on mRNA levels, apart from a greater effect of *atg-18* RNAi when compared to either *atg-4.1* and *atg-9* RNAi (Fig. S1B).

Comparing effects of RNAi on mRNA levels and lifespan, the only clear correspondence between the former and the latter involved *atg-2* and *atg-4.1* RNAi, which in N2 had no significant effect on either (Figure 1B, Fig. S1A; Table S2). However for the remaining 4 genes, there was no clear correspondence, though *atg-18* mRNA levels appear to be the lowest, in line with its greater effect on lifespan (Figure 1B-D, Fig. S1A; Table S2, Table S3). These results imply that RNAi efficacy and perhaps also gene-specific issues contribute to variation in *atg* RNAi effects on lifespan (discussed further below). While mRNA levels after RNAi under the various other conditions tested were not assayed, reduced IIS (including *daf-2(e1370)*) has been shown to intensify the RNAi response (Wang and Ruvkun, 2004), thus lack of effect on lifespan in *daf-2(e1370)* is unlikely to reflect suppression of mRNA knockdown.

### Effect of *atg* RNAi on *daf-2* Age is temperature dependent

Results of a previous study performed at 25°C appear to show no greater reduction in lifespan in *daf-2(e1370)* compared to *daf-2(+)* after autophagy knockdown, even when using the *atg-18(gk378)* deletion mutation (Toth et al., 2008) (Table S1), suggesting possible temperature dependence of *atg* RNAi effects. To explore this further, in parallel to tests at 20°C, we also compared effects of *atg* RNAi on lifespan in N2 and *daf-2(e1368)* at 25°C. Effects on *daf-2(e1370)* were not tested, partly because this mutant ceases feeding at 25°C (Gems et al., 1998) which would be expected to interfere with RNAi by feeding.

At 25°C, *atg* RNAi did not shorten N2 lifespan for any of the six genes tested, not even *atg-18* (summed data; Figure 1E, Table S2), i.e. culture at 25°C suppressed the life-shortening effect of *atg* RNAi in N2 (although in later tests at 25°C, life-shortening effects of *atg-13* and *atg-18* RNAi were seen; Table S9). Also notable is that the increases in lifespan with *atg-2* and *atg-13* RNAi at 25°C, described in our earlier study (Ezcurra et al., 2018), were not reproduced (discussed below).

At 25°C only *atg-18* RNAi significantly reduced *daf-2(e1368)* mean lifespan, by 15.5% (*p* < 0.0001) (summed data, Figure 1F, Table S2), a reduction that was significantly greater than in N2 (*p* = 0.0012, CPH, summed data only; Table S2). For survival curves comparing RNAi effects of individual *atg* genes on the two genotypes, see Fig. S2. In one case, *atg-9*, RNAi slightly increased *daf-2* lifespan (+9.8%, *p* = 0.011, summed data).

The initial tests showing that *bec-1* RNAi reduces *daf-2(e1370)* lifespan were performed at 15°C (Meléndez et al., 2003); moreover, a greater N2 life-shortening effect of *bec-1* RNAi at 15°C than 20°C has been reported (Chen et al., 2019). Taken together with the weaker RNAi effects at 25°C observed here, this suggested that stronger effects might be seen at 15°C. To explore this we compared effects of *bec-1* RNAi from L4 on N2 and *daf-2(e1370)* at 15°C and 20°C (*N* = 2). However, no suppression of *daf-2(e1370)* Age by *bec-1* RNAi was seen at either temperature (Figure 1G; Table S5), consistent with findings of an earlier study performed at 15°C (Hars et al., 2007) (Table S1). In these trials *bec-1* RNAi also modestly increased N2 lifespan at both temperatures (Table S5), surprisingly given that in previous trials (performed several years earlier) *bec-1* RNAi shortened N2 lifespan (Table S2). The reason for this discrepancy is unknown.

### FUDR can alter the effect of *atg* RNAi on lifespan

Since the 1980s FUDR, a thymidylate synthase inhibitor and anti-cancer drug, has sometimes been added to *C. elegans* survival trials to prevent progeny production (Gandhi et al., 1980; Mitchell et al., 1979). Notably, two previous reports that observed increases in *C. elegans* lifespan given *atg* RNAi employed FUDR. Our own study saw increases in N2 lifespan after *atg-2* and *atg-13* RNAi from L4 with 15 μM FUDR (Ezcurra et al., 2018). Another study that saw increases in N2 lifespan given adult-limited *atg-7*, *atg-9*, *bec-1* and *unc-51* RNAi used FUDR at a higher concentration, 800 μM (Hashimoto et al., 2009) (E. Nishida, personal communication).

To test for FUDR-dependent effects, we compared the impact of *atg* gene RNAi (*atg-2*, *atg-4.1*, *atg-9*, *atg-13*, *atg-18* and *bec-1*) on N2 lifespan at 20°C with 0, 15 or 800 μM FUDR (*N* = 2). With 0 or 15 μM FUDR, only shortening of lifespan was seen, and FUDR did not significantly alter the effects of RNAi (CPH, Figure 3A,B, Table S6).

**Figure 3.**
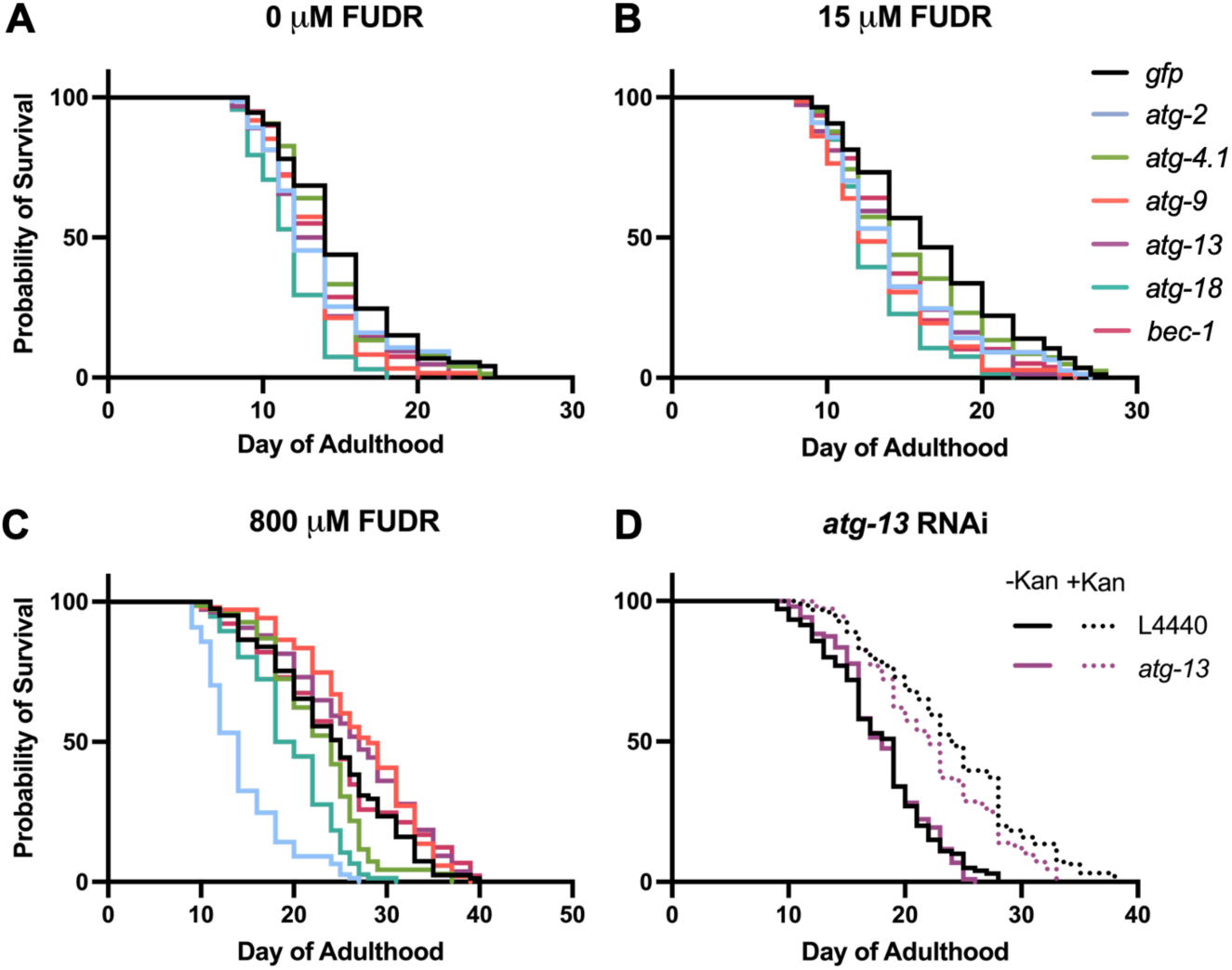
FUDR but not infection alters outcome on *atg* gene RNAi (20°C). (A-C) Effects of 0 mM, 15 μM and 800 μM FUDR. We note that in (A) the lifespan of the *gfp* RNAi control is somewhat lower than in other experiments (mean 14.96 days, Table S6); see Discussion for consideration of possible reasons for inter-trial variability. (D) No alteration by kanamycin of *atg-13* RNAi effect on N2 lifespan. Summed data, *N* ≥ 2; for individual trials and statistical comparisons, see Table S6, S7.

Addition of 800 μM FUDR increased the lifespan of the *gfp* RNAi negative control by 61.7% (*p* < 0.0001, Table S6), perhaps due to prevention of bacterial proliferation, which can otherwise shorten *C. elegans* lifespan (Garigan et al., 2002; Gems and Riddle, 2000). In the presence of 800 μM FUDR, *atg-9* and *atg-13* RNAi increased lifespan, by +12.5% (*p* = 0.014) and +8.7% (*p* = 0.031), respectively (summed data; Figure 3C, Table S6). Moreover, the life-shortening effect of *bec-1* RNAi, seen with 0 or 15 μM FUDR, was absent. Again, *atg-18* RNAi robustly reduced lifespan under all three conditions (Figure 3C, Table S6). The results using 800 μM FUDR are broadly in line with those of Hashimoto et al. (2009), where *atg-9* (and also *atg-7*, *bec-1* and *unc-51*) RNAi increased lifespan, and *atg-18* was one of only 2/14 genes tested where adult-limited RNAi decreased N2 lifespan. This suggests that the increases in lifespan after *atg* RNAi reported in that study could have reflected its use of 800 μM FUDR (Hashimoto et al., 2009).

### *atg* RNAi shortens lifespan in the absence of bacterial infection

Under standard laboratory culture conditions, *C. elegans* lifespan is limited by infection by the *E. coli* food source, such that prevention of bacterial proliferation substantially increases lifespan (Garigan et al., 2002; Gems and Riddle, 2000). The preceding results could imply that 800 μM FUDR suppresses life-shortening effects of *atg* RNAi by preventing bacterial infection. Xenophagy is generally protective against infection in *C. elegans* (Balla et al., 2019; Jia et al., 2009). Thus, reduction in lifespan given *atg* gene knockdown could reflect increased susceptibility to infection.

To probe this hypothesis, we compared effects on N2 lifespan of *atg-13* RNAi at 20°C in the absence or presence of the antibiotic kanamycin (Kan), to suppress bacterial infection. In the absence of Kan, *atg-13* RNAi did not alter lifespan in these trials (Figure 3D, Table S7), in contrast to the modest reductions in lifespan seen in other trials in this study (Table S2, S6) (for possible reasons for this discrepancy, see Discussion). Application of Kan extended *C. elegans* lifespan (+34.2%, *p* < 0.0001, summed data, Table S7), as previously observed (Garigan et al., 2002), and in its presence *atg-13* RNAi resulted in a modest decrease in lifespan (-6.7%, *p* = 0.0092, Figure 3D, Table S7). These results suggest that life-shortening effects of *atg* RNAi are not solely attributable to increased susceptibility to *E. coli* infection. Moreover, they do not indicate marked condition dependency in *atg* RNAi effects on lifespan with respect to *E. coli* proliferative status. We also conclude the increase in lifespan upon *atg-13* RNAi in the presence of 800 μM FUDR (Figure 3C) is not attributable to suppression of *E. coli* infection, but rather to some other, unidentified mechanism.

### *atg-18* RNAi robustly suppresses *glp-1(e2141)* Age

We next explored more widely the reproducibility and condition dependence of the requirement for autophagy of *C. elegans* longevity. Prevention of germline development by laser surgery or mutation increases lifespan in *C. elegans* hermaphrodites (Arantes-Oliveira et al., 2002; Hsin and Kenyon, 1999; Pires da Silva et al., 2024). Prior tests for possible autophagy dependence of such longevity have largely used the temperature-sensitive *glp-1(e2141)* <u>g</u>ermline <u>p</u>roliferation mutant, which is fertile at 15°C but sterile and with greatly reduced germline development at 25°C. A key study reported strong and reproducible suppression of *glp-1* Age by RNAi of five autophagy-related genes, including *atg-18* and *bec-1* (Lapierre et al., 2011) (previous findings summarized in Table S8).

We first tested the effect of *atg* RNAi on *glp-1* longevity with animals raised from L1 until L4 stage at 25°C, and maintained at 20°C thereafter, similar to previous studies (Table S8). Again, effects of *atg-2*, *atg-4.1, atg-9, atg-13*, *atg-18* and *bec-1* RNA were tested. In the RNAi control *glp-1* increased mean lifespan by +50.9% (*p* < 0.0001, summed data, Table S9). *glp-1* lifespan was significantly reduced by *atg-2*, *atg-4.1*, *atg-9*, *atg-18* and *bec-1* RNAi (but not *atg-13* RNAi), and effects were greater in *glp-1* than N2 in all cases except *atg-4.1* (Figure 4A, B, Figure 5, Table S9). However, suppression was only robust (of a large magnitude) for *atg-2* and *atg-18* RNAi (Figure 4B, Figure 5). Under all six *atg* RNAi conditions, *glp-1* still significantly increased mean lifespan, from +4.12% (*p* = 0.0003) for *atg-18* to +43.8% (*p* < 0.0001) for *atg-4.1* (summed data, 20°C, Table S9). This could imply either that *glp-1* Age is not fully autophagy dependent, or that it is but autophagy is not fully suppressed by the RNAi treatments used.

**Figure 4.**
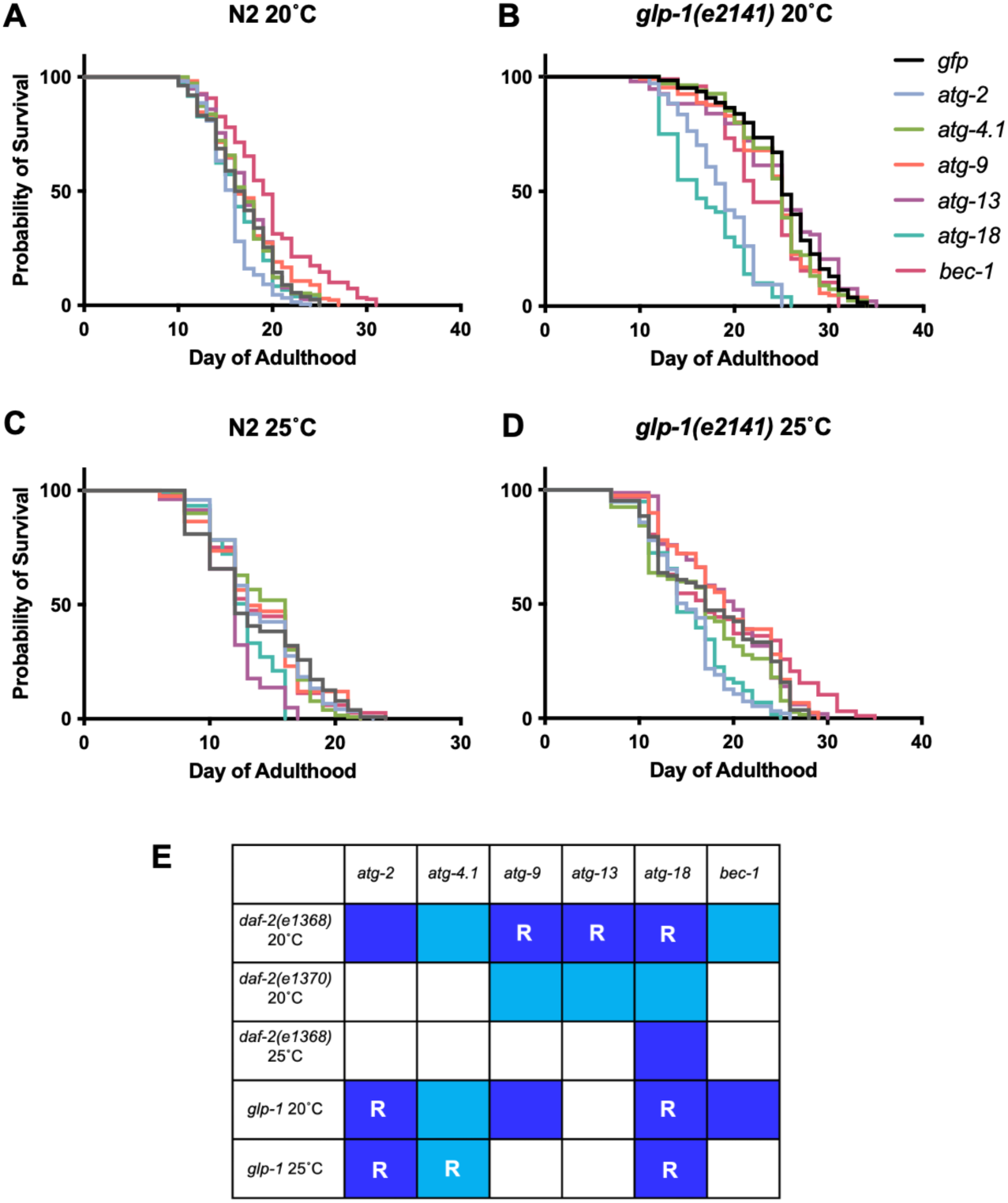
Degree of suppression of *glp-1(e2141)* longevity by *atg* gene RNAi differs greatly between genes. (A, B) 20°C. (A) Effects on N2. (B) Effects on *glp-1*. (C, D) 25°C. (C) Effects on N2. (D) Effects on *glp-1*. Summed data, *N* = 2-5; for individual trials and statistical comparisons, see Table S9. That *bec-1* RNAi increases N2 lifespan in (A) (+16.7%, *p* < 0.0001) but not (B) could imply an interaction with temperature during development, or merely variability of *atg* RNAi effects (see Discussion). (E) Overview of effects of suppression of Age by RNAi of genes specifying autophagy. Dark blue, reduction of *daf-2* or *glp-1* Age that is significantly greater than in N2. Light blue, reduction of *daf-2* or *glp-1* Age that is not significantly greater than in N2. R, robust suppression, i.e. knockdown reduces the extended lifespan of *daf-2* or *glp-1* to within <30% of the mean lifespan of N2 under the same RNAi. This designation (“robust”) indicates a high degree of suppression of the mutant longevity phenotype (see Figure 2, Figure 5 and Fig. S2). Note that the *bec-1* RNAi effect on *glp-1* at 20°C is not classified as robust here even though it reduces lifespan to within <30% of the mean lifespan of N2 under *bec-1* RNAi, since the fact that it does so partly reflects an increase in N2 lifespan, rather than a robust life-shortening effect on *glp-1*.

**Figure 5.**
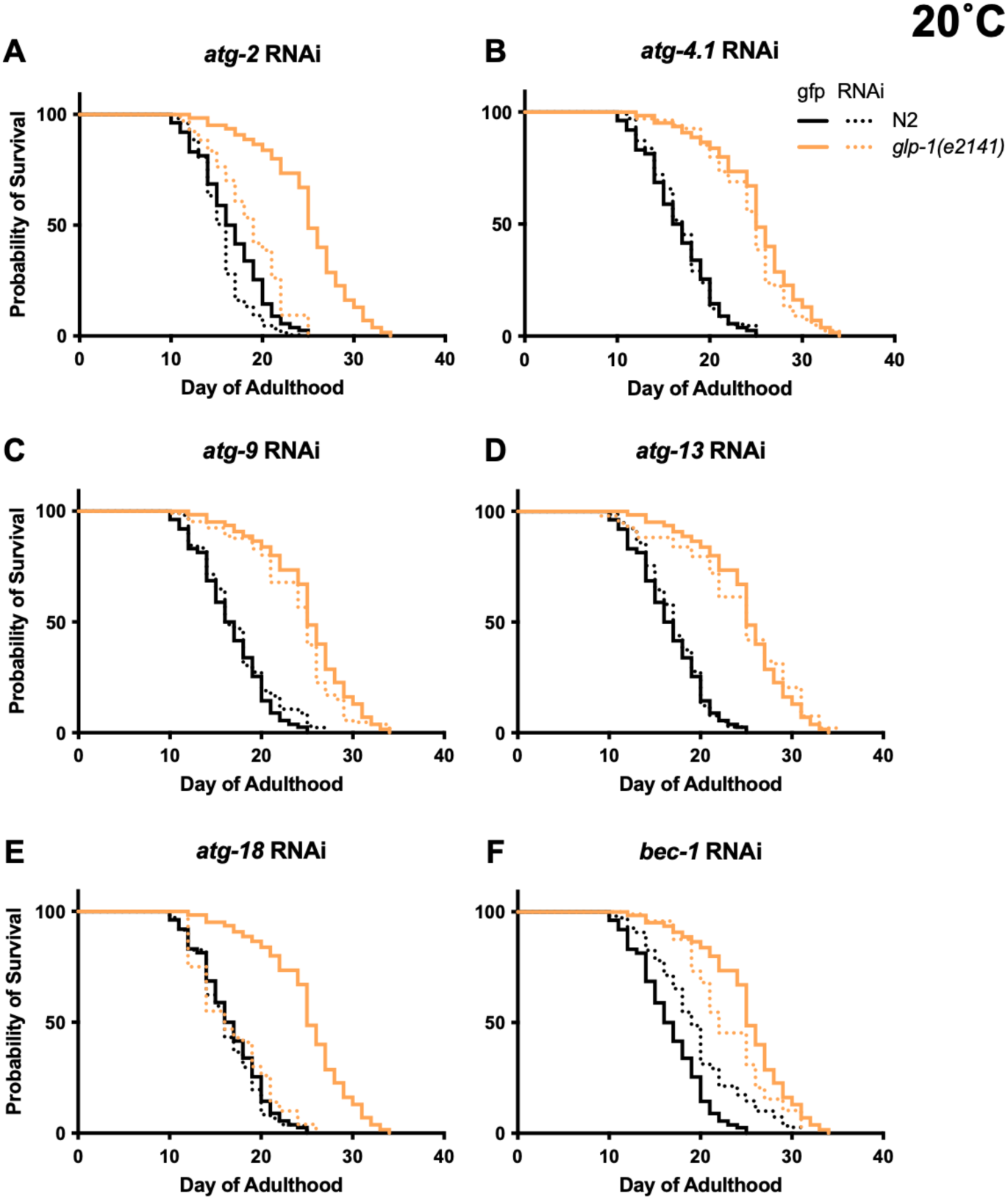
Degree of suppression of *glp-1(e2141)* longevity by *atg* gene RNAi differs greatly between genes (20°C), cf. Figure 4A,B (same data). Summed data, *N* = 2-5; for individual trials and statistical comparisons, see Table S9.

In tests with *daf-2(e1368)* life-shortening effects of *atg* RNAi were largely absent at 25°C (apart from *atg-18*) (Figure 1F, Table S2). We therefore wondered whether temperature might also influence the outcome of *atg* RNAi treatment in *glp-1* mutants. To test this we examined RNAi effects on lifespan at 25°C. Under these conditions, *glp-1* lifespan was significantly reduced by only *atg-2, atg-4.1*, and *atg-18* RNAi (Figure 4C, D, Fig. S3, Table S9), and effects were significantly greater in *glp-1* than N2 only with *atg-2* and *atg-18* RNAi (CPH analysis, Table S9). In summary, of the six genes tested only *atg-2* and *atg-18* RNAi robustly suppressed *glp-1* Age, and RNAi effects were weaker at 25°C.

For an overview of the effects of RNAi on *daf-2* and *glp-1*, see Figure 4E. In 11 of the 30 conditions tested RNAi knockdown of autophagy caused a greater reduction in lifespan in the long-lived mutant. In 8/30 the RNAi effect was robust, i.e. the mutant longevity was largely suppressed. This suppression was highly condition dependent, differing according to gene knocked down, temperature, *daf-2* allele used, and between *daf-2* and *glp-1*. Notably, only *atg-18* RNAi robustly suppressed longevity in both *daf-2* and *glp-1* mutants.

### Why are effects of *atg-18* RNAi stronger than those of other *atg* genes?

The particularly strong life-shortening effects of *atg-18* RNAi could reflect either a greater reduction in autophagy, or the presence in addition to its effects on autophagy of pleiotropic effects not directly related to autophagy. Given that longevity due to either *daf-2* mutation or germline loss are wholly dependent on the FOXO transcription factor DAF-16 (Hsin and Kenyon, 1999; Kenyon et al., 1993), we wondered whether *atg-18* RNAi might inhibit DAF-16. To probe this two approaches were taken. First, we used a constitutive dauer formation assay. High population density and food depletion causes *C. elegans* larvae to form developmentally arrested dauer larvae (Cassada and Russell, 1975). *daf-2* mutants undergo constitutive dauer arrest in a temperature-sensitive manner, and this is fully suppressed by *daf-16*(-) (Riddle et al., 1981). However, using a sensitive assay (*daf-2(m41)*, 22°C, giving a mix of dauer and non dauers), we detected no reduction in the number of dauers formed given *atg* gene RNAi (*atg-2*, *atg-13* and *atg-18*) (Fig. S4A).

Second, we tested a GFP reporter for a gene whose expression is elevated in *daf-2* mutants in a *daf-16*-dependent manner (*ftn-1*), using strains previously constructed (Ackerman and Gems, 2012). GFP levels were compared in *daf-2(m577)* and *daf-16(mgDf50)*; *daf-2* backgrounds. As expected, GFP levels were higher in *daf-2* than in *daf-16; daf-2* (Fig. S4B). RNAi did not suppress the *daf-2*-induced increase of *ftn-1::gfp* expression (Fig. S4B). These findings argue against a pleiotropic effect of *atg-18* on DAF-16 function.

### Inhibiting autophagy does not reduce vitellogenin accumulation

Finally, we further investigated the hypothesis that autophagy promotes biomass repurposing in *C. elegans*. Inhibition of yolk synthesis or of autophagy delays intestinal atrophy and yolk pool accumulation, suggesting that intestinal biomass is repurposed for yolk synthesis (Benedetto and Gems, 2019; Ezcurra et al., 2018; Sornda et al., 2019). In principle, this could involve repurposing into yolk protein (vitellogenin) or yolk lipid. To test the former possibility, wild-type hermaphrodites or *fog-2(q71)* (feminization <u>o</u>f <u>g</u>ermline) mutant females were subjected to *atg* RNAi. The *fog-2* mutant, which lacks self-sperm and so lays no eggs, was used to avoid possible effects of *atg* RNAi on fertility, reduction of which can increase vitellogenin levels within nematodes (Sornda et al., 2019).

For none of the six *atg* genes tested did RNAi detectably reduce yolk protein levels, either in N2 hermaphrodites or *fog-2* females (Figure 6). This implies that autophagic machinery, including that in the intestine, does not enhance yolk protein production. Thus, if intestinal biomass repurposing occurs, then it likely supports yolk secretion or yolk lipid production.

**Figure 6.**
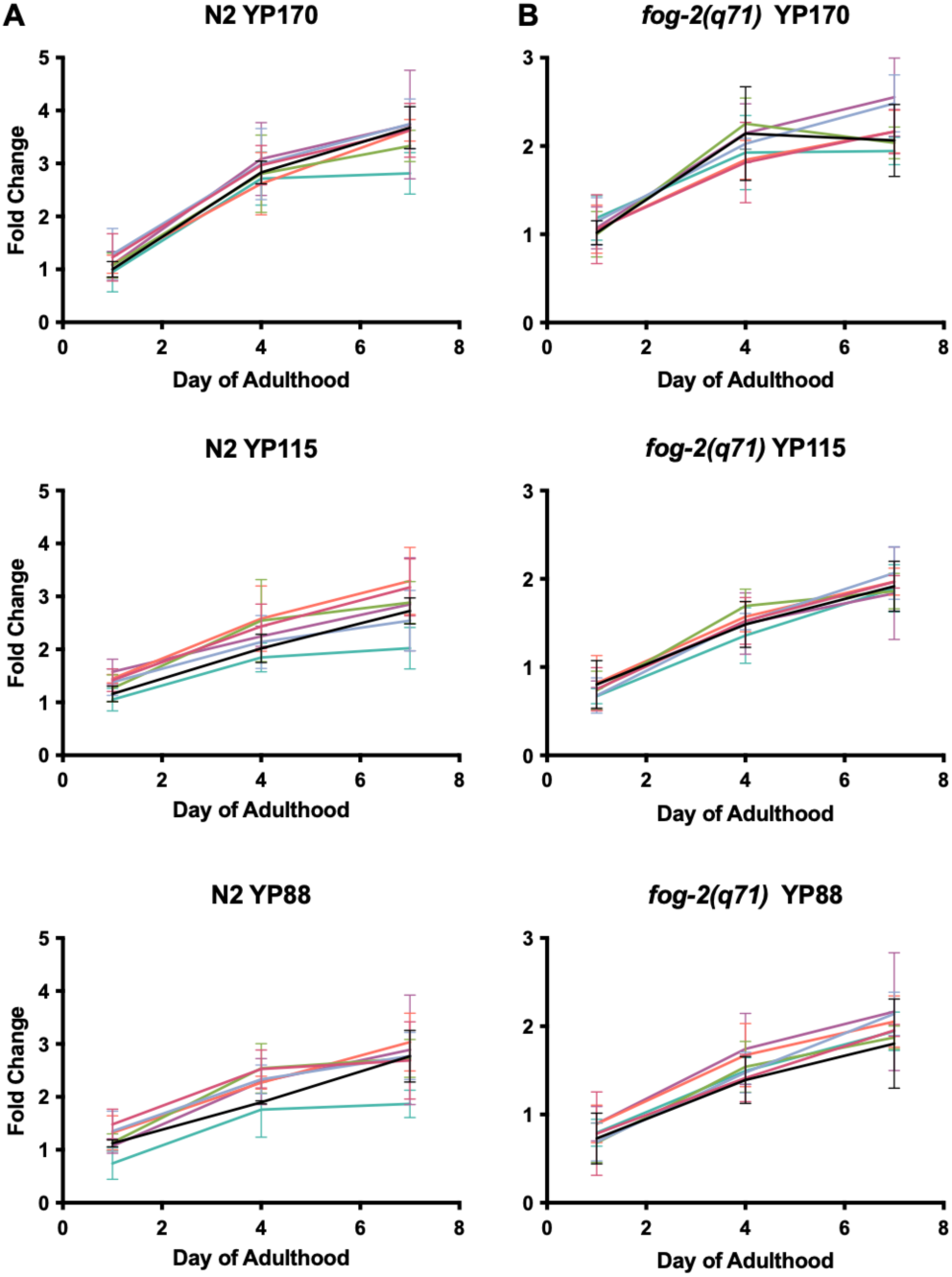
Absence of effect of *atg* RNAi on vitellogenin accumulation. Fold change of yolk proteins YP170, YP115 and YP88 in N2 and *fog-2* normalized to *gfp* day 1 (*N* = 3). In no case is vitellogenin level significantly different to that in the *gfp* RNAi control at any time point (two-way ANOVA, Table S10). For raw data for vitellogenin levels see Supplementary Dataset 3.

## Discussion

Overall, the effects of reducing *atg* gene function described here are ambiguous with respect to the role of autophagy in *daf-2* or *glp-1* longevity, neither strongly supporting or excluding it. However, they are consistent with a role of autophagy in the longevity of weaker, class 1 *daf-2* alleles at lower temperatures (Figure 1C,D,F,G, Table S2). Class 1 allele-limited suppression of *daf-2* Age has been seen previously, for example in epistasis tests with *daf-12* (encoding a dafachronic acid receptor) (Gems et al., 1998; Larsen et al., 1995) and *skn-1* (Nrf2-like transcription factor) (Tullet et al., 2008). This could reflect a role of autophagy in longevity assurance limited to conditions of mild IIS reduction, or sensitivity to differential effects of *daf-2* on distinct downstream signaling outputs, such as phosphatidylinositol 3-kinase and Ras signaling (Patel et al., 2008).

*atg* RNAi suppresses *daf-2(e1368)* Age at 20°C but not 25°C (Figure 1C,F, Table S2). Given that for hypomorphic *daf-2* alleles (such as *e1368* and *e1370*) many mutant traits, including Age, show some degree of temperature sensitivity (Gems et al., 1998; Riddle and Albert, 1997), the absence of suppression at 25°C may reflect the increased severity of the *daf-2(e1368)* mutant phenotype at this higher temperature. However, life-shortening effects of *atg* RNAi on N2 are also largely absent at 25°C, suggesting that additional mechanisms may also mediate such temperature sensitivity.

### Resolving discrepancies between past studies of autophagy dependence of *daf-2* Age

The increasing non-reproducibility of many experimental findings, that has been referred to as the *reproducibility crisis*, is a problem that particularly afflicts biological and biomedical research (Baker, 2016; Lithgow et al., 2017; Prinz et al., 2011; Ritchie, 2020; Voelkl et al., 2020). A strength of *C. elegans* as an experimental model is the relative ease with which such discrepancies can be resolved, at least in principle. This is thanks to the use of standardized culture conditions across the *C. elegans* research community, and of nematode strains based on the same isogenic wild-type strain (N2), plus the relatively low cost and short duration of experiments.

Regarding lifespan assays in particular, possible reasons for discrepant findings include clearly identifiable differences in experimental design, such as use or not of FUDR. Less obvious causes include cryptic variation in wild-type genetic background (Zhao et al., 2019), or subtle differences in culture conditions (e.g. due to batch variation in Bacto Peptone, a constituent of nematode growth medium) (Petrascheck, 2014).

Our findings potentially resolve several discrepancies between earlier studies relating to the possible role of autophagy in *daf-2* mutant longevity. Previous studies found that inhibiting autophagy either suppressed or enhanced *daf-2* Age, or had no effect (Table S1). One study showing suppression used the weak class 1 allele *daf-2(mu150)* (Hansen et al., 2008). This is consistent with our observation that, *atg-18* aside, *atg* RNAi can suppress Age in a class 1 but not a class 2 allele (Figure 1C,D Table S2).

Does *bec-1* RNAi suppress *daf-2(e1370)* Age? The initial test suggesting this was performed at 15°C, and employed RNAi by injection of the mothers of assayed individuals (Meléndez et al., 2003). In a subsequent study at 15°C, in which both mothers and adult progeny were exposed to *bec-1* RNAi by feeding, *daf-2(e1370)* lifespan appeared not to be shortened (Hars et al., 2007). Several further trials under different conditions did not observe a life-shortening effect either (Hashimoto et al., 2009; Toth et al., 2008). In the present study, we saw no shortening of *daf-2(e1370)* lifespan in summed data with adult-limited *bec-1* RNAi by feeding, at either 15°C or 20°C (Figure 1D,G, Fig. S3, Table S2, S5). Taken together with earlier evidence, our observations suggest that suppression of *daf-2(e1370)* Age by *bec-1* RNAi is not a readily reproducible finding.

Several studies reported lifespan extension following *atg* RNAi (Ezcurra et al., 2018; Hashimoto et al., 2009). Here we were able to reproduce this effect for several *atg* genes by applying high dose FUDR (800 μM), thus recapitulating findings by Hashimoto et al (2009), and potentially accounting for the life span increases seen in that study; it was only subsequent to that study that evidence emerged of the capacity of FUDR to alter effects of interventions that impact lifespan (Aitlhadj and Sturzenbaum, 2010; Anderson et al., 2016; Burnaevskiy et al., 2018; Van Raamsdonk and Hekimi, 2011; Zhao et al., 2019).

Returning to issues with replicating published findings: in an earlier study we observed increases in N2 lifespan given *atg-2* and *atg-13* RNAi (15 μM FUDR present) (Ezcurra et al., 2018). However, this finding proved not to be robust to replication (Figure 3B, Table S6), perhaps reflecting inherent effect variability, condition dependence of an unidentified kind, or a combination of the two. This again illustrates the value, as with the effects of *bec-1* RNAi on *daf-2(e1370)* Age, of repeated verification of effects of interventions on *C. elegans* lifespan.

Regarding the causes of variability between results of ostensibly identical tests performed under ostensibly identical conditions: one clue is provided by a study comparing results of lifespan assays performed across three sites under similar conditions. This revealed that inter-trial variation occurred mainly at each site over time, rather than between sites (Lucanic et al., 2017). One possibility is that this reflects batch variation in media components, such as the BactoPeptone constituent of nematode growth medium (Petrascheck, 2014).

As with *daf-2*, our tests with the *glp-1* germline proliferation mutant did not clearly support the view that mutant longevity is autophagy dependent. However, consistent with an earlier study (Lapierre et al., 2011), we observed that *atg-18* and *bec-1* RNAi reduce *glp-1* lifespan more than in wild type at 20°C.

### Why does *atg-18* RNAi more strongly suppress *daf-2* Age?

Of the six autophagy-determining genes tested here, RNAi of *atg-18* showed a greater capacity to suppress both *daf-2* and *glp-1* Age than the other five. *atg-18* appears to have replaced *bec-1* as the gene of choice for autophagy-related epistasis studies in *C. elegans* (Chang et al., 2017; Minnerly et al., 2017). Notably, *atg-18* was one of only 2/14 autophagy genes tested where RNAi during development caused a high level (>50%) of larval growth arrest or lethality (Hashimoto et al., 2009). The more marked effects of *atg-18* RNAi could reflect either greater inhibition of autophagy, or pleiotropy in which processes other than autophagy are altered.

Regarding pleiotropy, several proteins in the canonical autophagy pathway have recently been found to participate in other processes. For example, ATG8 (LGG-1 in *C. elegans*) functions in trafficking of single-membrane organelles (Nieto-Torres et al., 2021), and *C. elegans* ATG-16.2 contributes to neuronal exopher formation via its WD40 domain (Yang et al., 2024). In an as yet unexplained instance of *atg* gene idiosyncrasy, mutation of *atg-16.2*, *atg-18* or *bec-1* retards the cell cycle in *C. elegans* germline cells, while that of *atg-7* does not (Ames et al., 2017).

Concerning *atg-18*, several previous studies suggest possible pleiotropy. ATG-18 is a predicted WIPI (WD repeat protein Interacting with PhosphoInositides) family member. In humans there are four WIPI proteins, WIPI1 - WIPI4. *C. elegans atg-18* is more closely related to WIPI1/WIPI2, while *epg-6* (Ectopic P Granules 6) more closely resembles WIPI3/WIPI4 (Lu et al., 2011). Notably, *atg-18* and *epg-6* deletion mutations appear to cause similar reductions in levels of autophagy, yet only *atg-18* shortens lifespan (Takacs et al., 2019).

Next, *C. elegans* studies of rescue of longevity and fat storage in *daf-2(e1370); atg-18(gk378)* double mutants by tissue-specific expression of *atg-18*(+) revealed a major role of *atg-18* in food-sensing chemosensory neurons, potentially reflecting a vesicle trafficking role in neurosecretion (possibly of neuropeptides) (Jia et al., 2019; Minnerly et al., 2017). Whether such a role is unique to *atg-18* or shared by other *atg* genes remains unclear.

Third, recent evidence suggests that ATG-18 activates HLH-30/TFEB (Helix Loop Helix/Transcription Factor EB). *C. elegans* HLH-30 is a master transcriptional regulator of autophagy and lysosomal biogenesis, that becomes nuclear localized in *daf-2(e1370)* and *glp-1(e2141)* mutants, and whose inhibition reduces longevity in both contexts (Lapierre et al., 2013; Lin et al., 2018; Wong et al., 2023). Notably, among both genes up-regulated upon over-expression of *atg-18*, and genes down-regulated in an *atg-18(gk378)* mutant, ones with HLH-30 elements in their promoters are over-represented (Schmauck-Medina et al., 2026). Also, heat shock-induced nuclear localization of HLH-30 is suppressed by *atg-18* RNAi, while longevity resulting from *atg-18* over-expression is suppressed by *hlh-30* RNAi.

Inhibition of HLH-30 by *atg-18* RNAi could potentially explain its unusually strong life-shortening effects. If correct, this could reflect greater suppression of autophagy, but also, in principle, other effects of HLH-30 inhibition. HLH-30 affects a variety of traits, including adult reproductive diapause (Gerisch et al., 2020), proteostasis (Shalash et al., 2025), sex differences in immunity (Sohn et al., 2025), and resistance to various insults, including infectious pathogens (El-Houjeiri et al., 2019; Visvikis et al., 2014; Wani et al., 2021), starvation (Harvald et al., 2017; O’Rourke and Ruvkun, 2013; Settembre et al., 2013), and oxidative and heat stress (Lin et al., 2018).

While in at least some of these cases autophagy likely contributes to *hlh-30*-mediated effects, given that this transcription factor also promotes expression of many genes unrelated to autophagy (Chen et al., 2017; Lin et al., 2018; Shalash et al., 2025; Sohn et al., 2025; Visvikis et al., 2014), other functions may also be involved. For example, in response to infection HLH-30 can activate expression of signaling (including IIS), autophagy-related and immunity-related genes, and both of latter can contribute to infection resistance (Chen et al., 2017; Sohn et al., 2025; Visvikis et al., 2014). In conclusion, if the more severe effects of *atg-18* RNAi on lifespan reported here are attributable to reduced HLH-30 activity, this could well reflect greater reduction of autophagy. However, it remains possible that other functions controlled by HLH-30 also play a role, e.g. relating to immunity (Chen et al., 2017; Visvikis et al., 2014) and protein quality control (Shalash et al., 2025). Also notable is that HLH-30 regulates gene expression combinatorially with DAF-16 (Lin et al., 2018), mutation of which fully suppresses *daf-2(rf)* longevity (Kenyon et al., 1993).

Finally, recent findings suggest that dependence of *glp-1* Age on *atg-18* is in part unrelated to its role in autophagy, but instead to an alternative pathway involving PCK-2 phosphoenolpyruvate carboxykinase (Shioda et al., 2026). Taken together, emerging findings argue that one may not safely conclude that effects of *atg-18* on lifespan are attributable solely to its role in autophagy.

### Autophagy in biomass repurposing during programmatic aging

Autophagic processes play a major role in tissue atrophy and degeneration related to biomass repurposing in semelparous organisms (that reproduce once and then die), particularly plants (Avila-Ospina et al., 2014; Gems et al., 2021). Previous studies support the hypothesis that intestinal biomass is repurposed for synthesis of yolk that, subsequent to egg laying, is vented to support larval development (Kern et al., 2021), and that autophagy facilitates this biomass conversion (Ezcurra et al., 2018; Sornda et al., 2019). If late-life mortality is promoted by intestinal atrophy, then preventing it should extend lifespan. Consistent with this, intestinal atrophy and yolk production are suppressed in *daf-2* mutants (Depina et al., 2011; Ezcurra et al., 2018), and blocking yolk production both retards intestinal atrophy and modestly increases lifespan (Ezcurra et al., 2018; Murphy et al., 2003; Sornda et al., 2019), in part by increasing resistance to late-life *E. coli* infection (Wang et al., 2026). This could imply that inhibiting autophagy, by retarding intestinal atrophy, could extend lifespan under some conditions, and several earlier studies observed life extension after *atg* RNAi (Ezcurra et al., 2018; Hashimoto et al., 2009; Wilhelm et al., 2017).

The results of the present study are in line with the view that intestinal atrophy, though a salient feature of *C. elegans* senescence, by itself contributes only weakly to late-life mortality. Consistent with this, inhibition of *daf-2* using auxin-induced degradation of DAF-2 protein can strongly increasing lifespan even when initiated only at advanced ages, after intestinal atrophy has occurred, and without any detectable reversal of major senescent pathologies (Molière et al., 2024; Venz et al., 2021). One possibility is that *atg* RNAi has antagonistic effects on late-life mortality: modestly reducing it due to suppression of intestinal atrophy (as seen when vitellogenin synthesis is inhibited) (Ezcurra et al., 2018; Murphy et al., 2003; Sornda et al., 2019), but also increasing it, due to disruption of essential cellular functions.

The gut-to-yolk biomass repurposing hypothesis drew particularly on the observation that blocking yolk protein (vitellogenin) synthesis or autophagy reduced intestinal atrophy and pseudocoelomic yolk pool size (Ezcurra et al., 2018; Sornda et al., 2019); given that yolk pools contain yolk protein and lipid (Ezcurra et al., 2018; Garigan et al., 2002; Herndon et al., 2002; Palikaras et al., 2017), such putative biomass repurposing could increase production of yolk protein, yolk lipid or both. Here we tested whether *atg* RNAi reduces yolk protein levels, and found that it did not (Figure 6). If the repurposing hypothesis is correct, then this would support the view that promotion by autophagy of repurposing of intestinal biomass into yolk involves yolk lipid rather than yolk protein, or overall yolk secretion. In principle, this could involve mass export of stored lipid, either by means of lipophagy, or perhaps secretory autophagy (Ponpuak et al., 2015). As an aside, we note that there is evidence that vitellogenin levels can affect autophagy (Seah et al., 2016).

### Concluding remarks

This study reassesses the claim that *daf-2* Age is autophagy dependent. While the results do not entirely prove or disprove this claim, they clearly show that evidence of dependence varies greatly with context, and suggest that autophagy only contributes substantially to *daf-2* Age given a weak reduction in IIS.

One limitation in establishing the role of autophagy in *daf-2* mutant longevity is the difficulty of reliably measuring its level of activity. While several assays using fluorescent reporters to identify putative autophagosomes suggest possible increases in autophagy levels (Chang et al., 2017; Hansen et al., 2008; Meléndez et al., 2003), *daf-2* mutants also show marked reductions in both protein synthesis rate (Depuydt et al., 2013) and turnover rate (Dhondt et al., 2016; Visscher et al., 2016), including reduced ATG-18 turnover rate (Visscher et al., 2016). This is more consistent with a reduction rather than an increase in autophagy, and in line with observed suppression of intestinal atrophy in *daf-2* mutants (Ezcurra et al., 2018).

One issue raised by condition dependence of a given effect is that it creates a risk of condition selection bias, where choice of experimental conditions can bias the outcome of an experiment. Where condition dependence has been identified, condition selection bias may be avoided by selection of multiple conditions to test a given hypothesis. In the present case, this could entail comparing effects on *daf-2(e1368)* and *daf-2(e1370)* at 20°C, ideally without FUDR use, and RNAi tests with multiple *atg* genes, but perhaps not *atg-18*, at least until the reason for its idiosyncratic effects are understood. Another factor to take into account is the apparent tendency of results of *C. elegans* lifespan assays to vary over time (Lucanic et al., 2017). General questions relating to how to deal with condition dependency have been discussed previously (Munafò and Davey Smith, 2018; Voelkl et al., 2020).

A second, wider issue is that condition dependence risks generating a situation where different research articles make conflicting claims, supporting different views held by different groups of researchers. For science to progress effectively it is necessary for research communities to resolve discrepancies between published findings, though the work required can be tedious, a task akin to washing the dishes in a communal household. It requires identifying and flagging findings that are either condition dependent or, seemingly, unreproducible and, where possible, distinguishing the two. For *C. elegans* at least, such a tidying process is highly feasible, as is use of methodologies devised to reduce such discrepancies (Driscoll et al., 2025; Lucanic et al., 2017; Petrascheck and Miller, 2017), which is a particular virtue of this model organism. More widely, condition dependency and conditional selection bias risk diminishing the reliability of research findings in many scientific disciplines.

## Materials and Methods

### Culture methods and strains

*C. elegans* maintenance was performed using standard protocols (Brenner, 1974). Unless otherwise stated, all strains were on nematode growth media (NGM, containing Bacto Peptone) with plates seeded with *E. coli* OP50 to provide a food source. An N2 hermaphrodite stock recently obtained from the Caenorhabditis Genetics Center was used as wild type (N2H) (Zhao et al., 2019). Genotypes of most mutants used are as described in Wormbase (www.wormbase.org). Strains used included CB4027 *glp-1(e2141)*, GA633 *daf-2(m577)*; *wuIs177 [Pftn-1::gfp lin-15(+)]*, GA643 *daf-16(mgDf50)*; *daf-2(m577)*; *wuIs177 [Pftn-1::gfp lin-15(+)]*], GA1930 *daf-2(e1370)*, GA1945 *daf-2(m41)*, GA1960 *daf-2(e1368)*, and JK574 *fog-2(q71)*.

### Constitutive dauer larva formation assay

*daf-2(m41)* animals were maintained on RNAi feeding strains for at least two generations prior to analysis. For the dauer larva formation assay, performed at 22°C, 10 L4-stage hermaphrodites were transferred onto RNAi plates and allowed to lay eggs for approximately 6 hr, after which adults were removed. Dauers were scored at 166 hr.

### Epifluorescence microscopy

Nematodes were anaesthetized with 10 μl 2 mM levamisole on 2% agar pads prior to imaging. For imaging, we used either a Zeiss Axioskop 2 plus microscope with Hamamatsu ORCA-ER digital camera C4742-95 and Volocity 6.3 software (Macintosh version) for image acquisition; or an ApoTome.2 Zeiss microscope with a Hamamatsu digital camera C13440 ORCA-Flash4.0 V3 and Zen software for image acquisition.

### RNA-mediated interference (RNAi)

RNAi by feeding was performed using RNAi plasmids transformed into *E. coli* OP50(xu363) as previously described (Xiao et al., 2015). Bacterial transformants were selected on LB agar plates containing 10 µg/ml tetracycline and 25 µg/ml carbenicillin. Inserts of all RNAi feeding clones were confirmed by sequencing. Origins of plasmids in RNAi feeding strains: *atg-13*, *bec-1*: Ahringer library (Kamath et al., 2003); *atg-2*, *atg-4.1*, *atg-9*: Vidal library (Rual et al., 2004). As an *atg-18* RNAi feeding clone was not available in local RNAi libraries, a custom *atg-18* RNAi plasmid was generated by cloning the target sequence into the L4440 vector using primers gcctccacttcctgttgaag and gagactcttttcgtcggca.

For RNAi induction, bacteria were grown for 16 h at 37°C with shaking (200 rpm) in 5 ml LB supplemented with 25 µg/ml carbenicillin. Overnight cultures were diluted 1:100 into fresh LB containing 25 µg/ml carbenicillin and grown for a further 4 h or until reaching an OD₆₀₀ of ∼0.4. Expression of double-stranded RNA was induced by addition of 1 mM IPTG, and cultures were incubated for an additional 1 h. Following induction, cultures were allowed to cool to room temperature, concentrated five-fold by centrifugation, and seeded onto NGM plates supplemented with 25 µg/ml carbenicillin and 1 mM IPTG. Seeded plates were allowed to dry at room temperature before use.

To minimize IPTG degradation, RNAi plates were stored in foil-lined boxes. Typically, unseeded plates were stored at 4°C for up to one month, while seeded plates were used within two weeks. RNAi treatment was initiated from the L4 larval stage unless otherwise stated.

## Survival analysis

Nematodes were maintained at a density of 25-30 per plate, and transferred daily during the egg laying period, and every 6-7 days thereafter. The L4 stage was defined as day 0. Mortality was scored every 1-2 days, with worms scored as alive if they showed any movement, either spontaneously or in response to gentle touch with a worm pick.

### RNA extraction, cDNA synthesis and RT-qPCR

Approximately 100–200 day 4 adult animals per treatment group were lysed in 250 µl of TRIzol™ reagent by vortexing for 10 min at 4°C, followed by a 10 min incubation on ice; this cycle was repeated three times. RNA was purified using the RNeasy Mini Kit (QIAGEN, Cat. No. 74104) according to the manufacturer’s instructions, including on-column DNase digestion using the RNase-Free DNase Set (QIAGEN, Cat. No. 79256). cDNA was synthesized from 100 ng of total RNA using the SuperScript™ First-Strand Synthesis System (Thermo Fisher Scientific, Cat. No. 10684803).

PCR primers were designed using Primer-BLAST with the following parameters: maximum primer length of 20 nucleotides, PCR product size of 70–200 bp, and exon–exon spanning where possible. Primer efficiency and R² values were determined from standard curves to ensure suitability for quantitative analysis and comparable amplification efficiencies between primer pairs. Primers were used at a final concentration of 2 µM.

Quantitative PCR was performed using Fast SYBR™ Green chemistry on a QuantStudio™ 6 Flex Real-Time PCR System (Applied Biosystems) with 384-well plates and a total reaction volume of 10 µl. Cycling conditions were 95°C for 20 s, followed by 40 cycles of 95°C for 1 s and 65°C for 20 s, with fluorescence data acquisition during the annealing/extension step. Melt curve analysis was performed following amplification to confirm product specificity.

A full list of primers used in this study is provided in Table S11. mRNA levels were normalized to mRNA levels from two housekeeping genes, *cdc-42* and *pmp-3* (Hoogewijs et al., 2008).

### Electrophoresis of *C. elegans* yolk protein

N2 and *fog-2(q71)* worms were synchronized by performing an egg lay and allowing nematodes to grow at 20°C until they reached the L4 stage. L4 worms were then transferred to fresh seeded RNAi plates and maintained under standard culture conditions at 20°C. Five worms were harvested at day 1, day 4 and day 7 into microcentrifuge Eppendorf tubes containing 10 μl M9 buffer. Before running the gel, 10 μl of 2x Laemmli sample buffer was added. Samples were incubated at 70°C and vortexed periodically for 15 min. Samples were then incubated at 95°C and vortexed periodically for 5 min, and were centrifuged at 6,000 rpm for 15 min.

10 μl of sample was loaded into wells of Invitrogen NuPAGE Bis-Tris protein gels. 5 μl of CozyHi pre-stained protein ladder was loaded at the left side of the gels. The running buffer used was 5% XT MOPS (Bio-Rad). The gels were then run at 150V for 2 hr. Gels were removed from cassette and placed in 100 ml of Coomassie staining solution overnight. Gels were then washed with distilled water three times, then placed into destaining solution and soaked for 40 min. Gels were then washed with distilled water and stored at 4°C until imaged. Gels were imaged and saved as 8 bit grayscale TIF files. Images of gels were analysed using Fiji software. Bands of interest (e.g. myosin, YP170, YP115, YP88) were selected manually based on their molecular weight, and intensity of the bands was measured and exported to Microsoft Excel for further analysis.

## Data analysis and statistics

Data were plotted using ePrism 9.0 (GraphPad Software, Boston, MA, USA) or Jupyter Notebook using Python with the matplotlib, pandas and NumPy libraries. Statistical tests were performed on raw data using either Prism 9 or JMP Pro 15 (JMP Statistical Discovery LLC, Cary, NC, USA) unless otherwise stated. The specific tests and post hoc corrections performed are described in the figure legends. To detect alterations in lifespan, the log rank test was used. To compare the magnitude of changes in lifespan, Cox Proportional Hazard (CPH) analysis was used. RT-qPCR data was analyzed using the ΔΔCt method and t-tests. To compare yolk protein levels, a two-way ANOVA was used. For lifespan trials, no statistical methods were used to predetermine sample size. The experiments were not randomized. The investigators were not blinded to allocation during experiments and outcome assessment.

## Acknowledgments

We thank Georg Fuellen (Rostock University), Kailiang Jia (Florida Atlantic University), Alicia Meléndez (Queens College-CUNY), Eisuke Nishida (RIKEN), and John Labbadia, Om Patange and Hongyuan Wang (UCL) for useful discussion, and/or comments on the manuscript, and Minh Tran Dang, Anna Girtle, Changtai Li, Gadea Meecham-Garcia and Suzie Mishima for minor research contributions. Some strains were provided by the Caenorhabditis Genetics Center, which is funded by NIH Office of Research Infrastructure Programs (P40 OD010440).

## Data availability statement

The datasets used and/or analyzed during the current study are available with this article, and also from the corresponding author David Gems on request.

## Author contributions

David Gems, Conceptualization, Funding acquisition, Methodology, Project administration, Writing - original draft, Writing - review and editing; Hannah Chapman, Kuei Ching Hsiung, Investigation, Methodology, Writing - review and editing. Isadora Rawlinson, Xiaoyu Sun, Xiaoya Wei, Investigation.

## Additional files

Supplementary information. Supplementary Tables 2-7, 9-11. Supplementary Dataset 1, 2, 3.

