## Supplementary information for "Effects of knockdown of autophagy pathway genes on *C. elegans* longevity are highly condition dependent"

Contents Summary

**Supplementary Figure 1.** Efficiency and relative strength of *atg* gene knockdown by RNAi plasmid vectors (N2).

**Supplementary Figure 2.** Degree of suppression of *daf-2(e1368)* longevity by *atg* gene RNAi differs greatly between genes (25˚C).

**Supplementary Figure 3.** Degree of suppression of *glp-1(e2141)* longevity by *atg* gene RNAi differs greatly between genes (25˚C).

**Supplementary Figure 4.** No evidence of *atg* RNAi effects on *daf-16*.

**Supplementary Table 1.** Previous reports of effects of inhibition of autophagy on *daf-2* Age.

**Supplementary Table 8.** Previous reports of effects of inhibition of autophagy on GSC(-) Age.

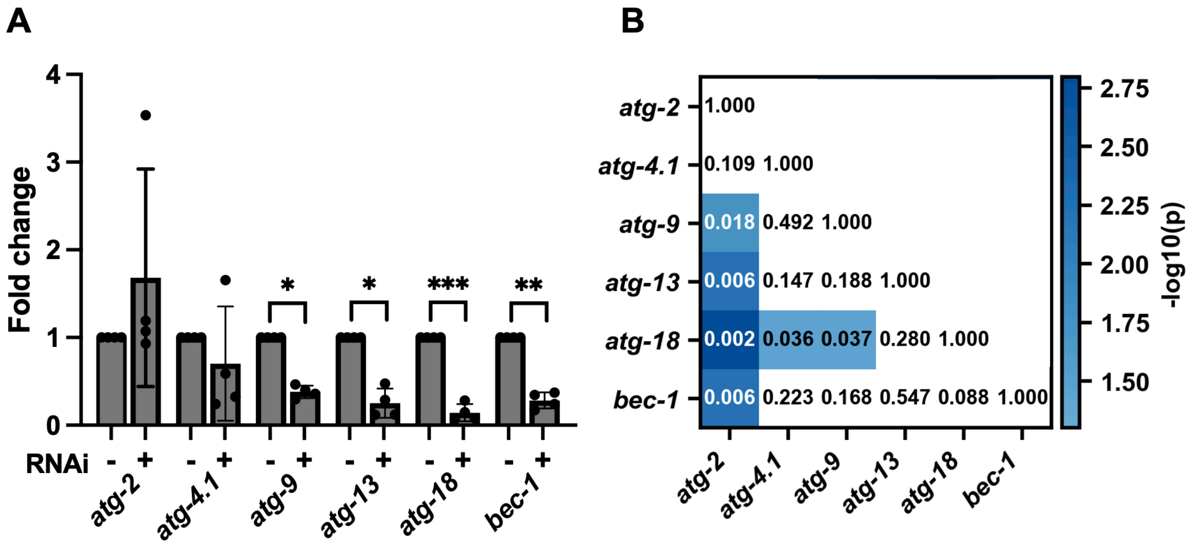

**Supplementary Fig. 1**. Efficiency and relative strength of *atg* gene knockdown by RNAi plasmid vectors (N2). (A) Relative *atg* mRNA levels following gene-specific RNAi treatment, calculated from ΔCt values relative to a *gfp* RNAi control. Fold change in expression was calculated as 2^−ΔΔCt. Statistical significance was assessed for each gene by unpaired t-test comparing ΔCt values between control and RNAi-treated conditions. Data are shown as mean ± SEM (4 independent RNAi replicates), with individual data points indicated. (B) Pairwise comparison of knockdown efficiency between *atg* genes. Statistical significance was assessed by pairwise comparison of ΔΔCt values across four independent trials using unpaired t-tests (Welch's correction applied where variances differed). Values shown within the heatmap indicate raw p-values, while color intensity represents –log10(p).

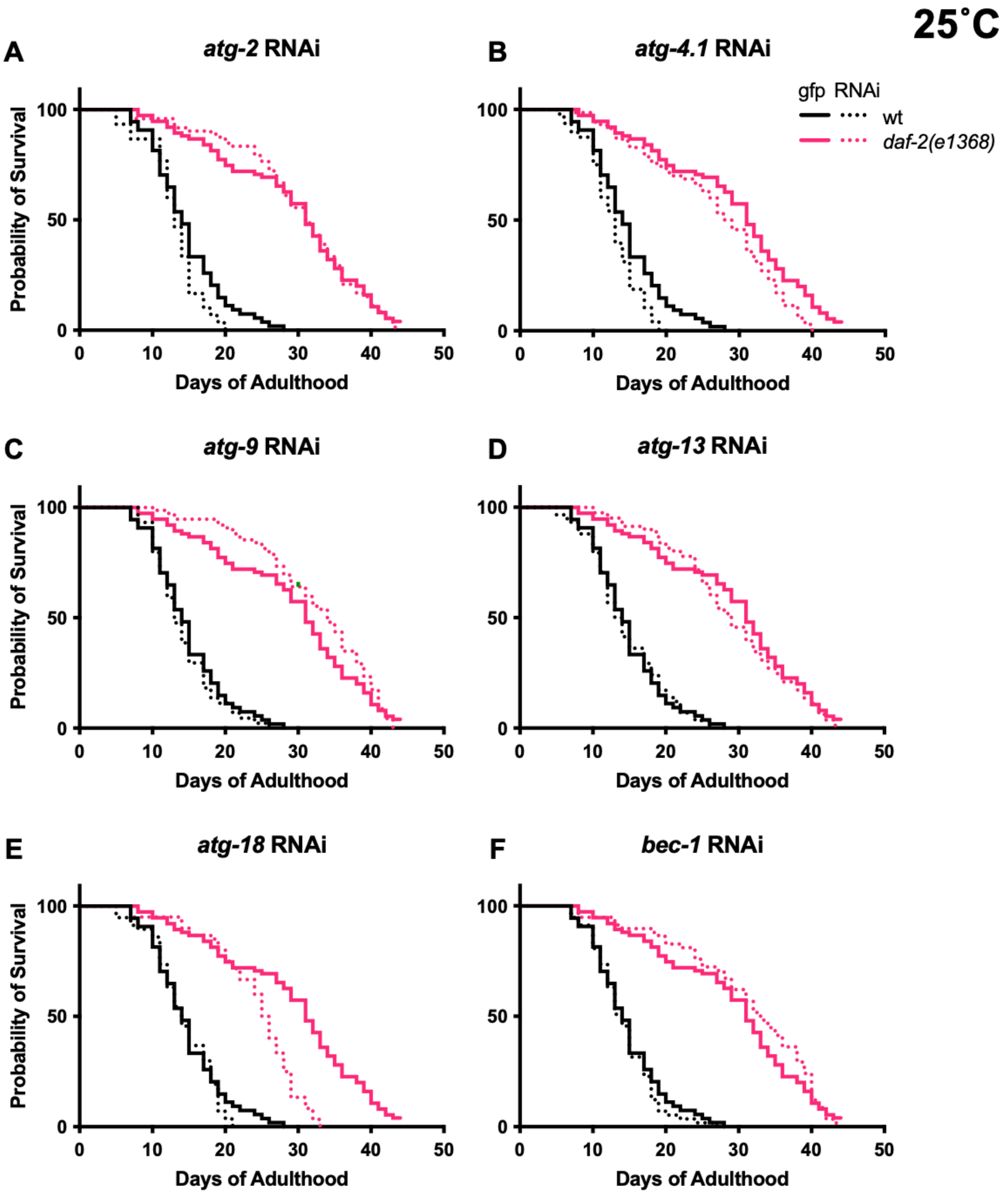

**Supplementary Fig. 2.** Degree of suppression of *daf-2(e1368)* longevity by *atg* gene RNAi differs greatly between genes (25˚C). Summed data, *N* = 2; for individual trials and statistical comparisons, see Table S2.

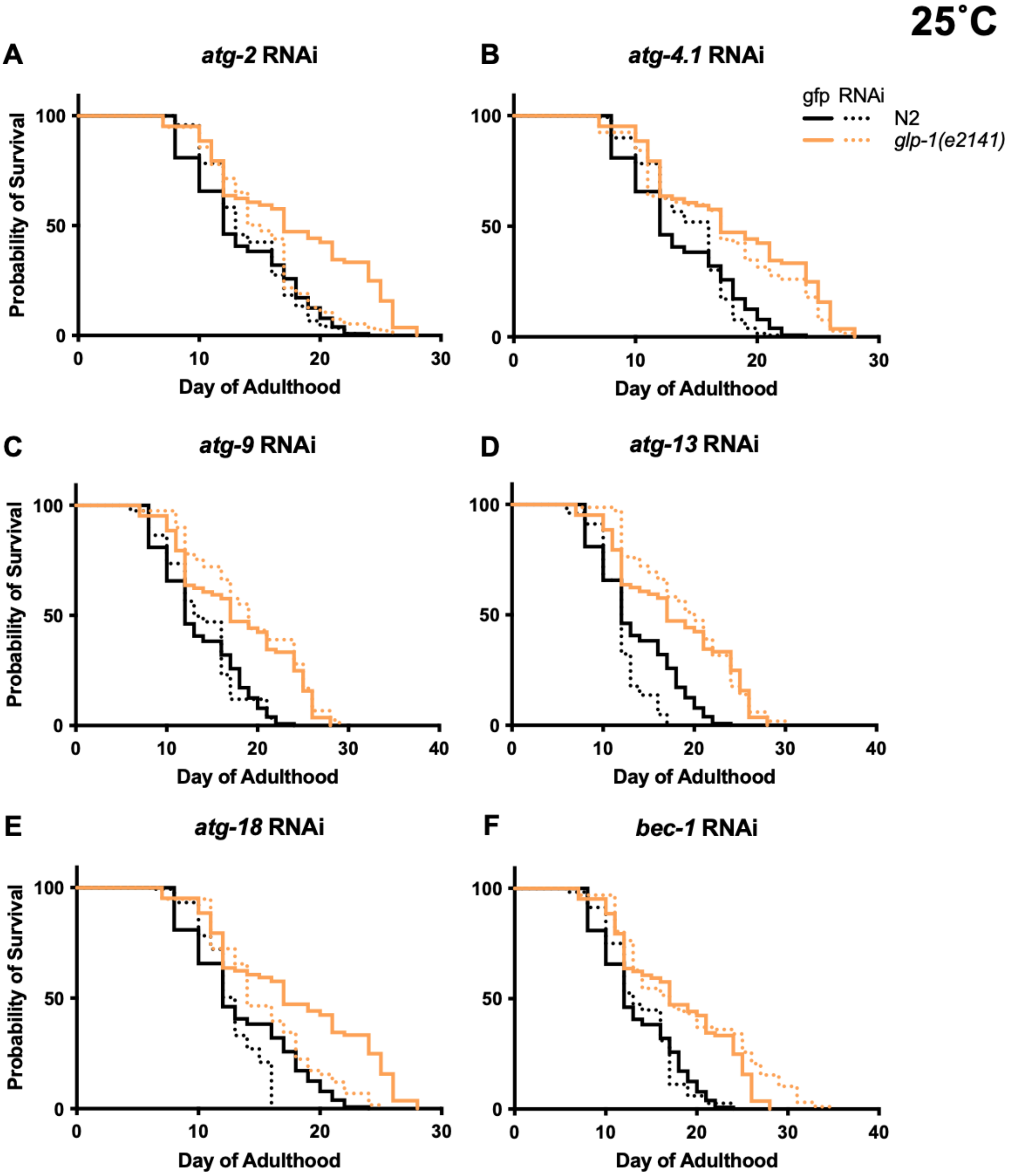

**Supplementary Fig. 3.** Degree of suppression of *glp-1(e2141)* longevity by *atg* gene RNAi differs greatly between genes (25˚C). Summed data, *N* = 2; for individual trials and statistical comparisons, see Table S9.

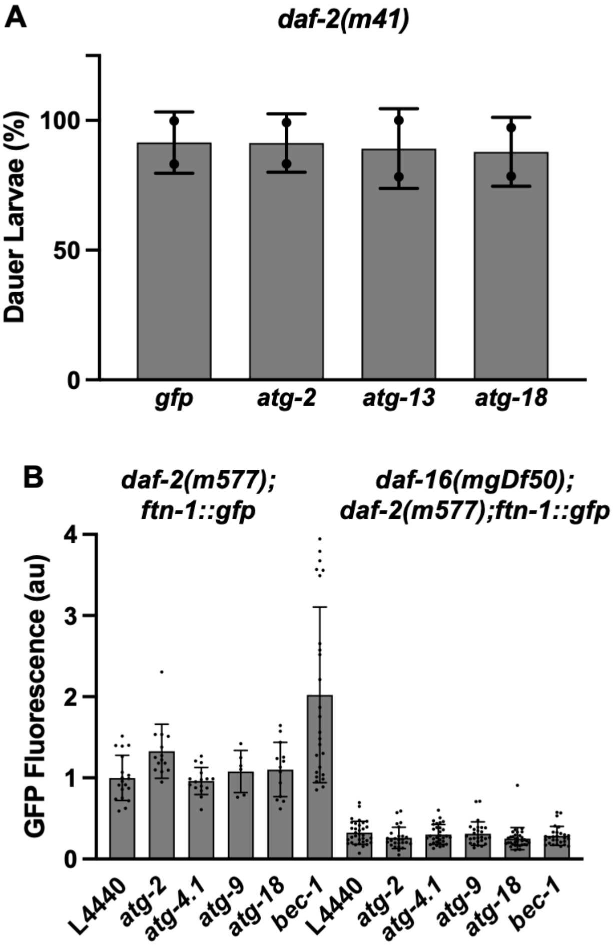

**Supplementary Fig. 4**. No evidence of *atg* RNAi effects on *daf-16*. Daf-c suppression assay was employed, using the class 1 mutant *daf-2(m41)* at 22.5˚C, previously found to form ~75% dauers (Gems et al., 1998). Larval progeny of animals maintained for two generations on RNAi were examined. (B) No effects of *atg* RNAi on *daf-16* target gene expression. The reporter tested was based on the gene *ftn-1* (ferritin) (Ackerman and Gems, 2012).

### Supplementary Table 1: Previous reports of effects of inhibition of autophagy on *daf-2* Age

| *C. elegans* genotype | *atg* gene | Mammalian gene | Form of gene inhibition | Age at initiation of RNAi | Temp | FUDR concentration | % change in *daf-2* mean LS cf RNAi or mutant control (*p*, log rank)[1] | % change in *daf-2(+)* control mean LS cf RNAi control (*p*, log rank)[1] | Greater reduction in LS in *daf-2*? (Cox proportional hazard) | *N*[2] | Source |
|---|---|---|---|---|---|---|---|---|---|---|---|
| *daf-2(e1370)* | *bec-1* | *BECN1* | RNAi by microinjection | Injection of previous generation | 15˚C | 0 | -46.6 (median lifespan) ($p<0.001$) | -14.2 (median lifespan) (no statistics) | Yes ($p<0.001$) | NS | (Meléndez et al., 2003) |
| *daf-2(e1370)* | *atg-7* | *ATG7* | RNAi by feeding | Young adult | 15˚C | 0 | Slight reduction (data not shown) | No effect (data not shown) | Not reported | NS | (Hars et al., 2007) |
| *daf-2(e1370)* | *atg-7* | *ATG7* | RNAi by feeding | Young adult of previous generation (RNAi during development) | 15˚C | 0 | -27.7 ($p$=5.4 x $10^{-9}$) | -28.5 ($p$=4.7 x $10^{-9}$) | Effect "appeared similar" (no statistical test) | NS | (Hars et al., 2007) |
| *daf-2(e1370)* | *bec-1* | *BECN1* | RNAi by feeding | Young adult of previous generation (RNAi during development) | 15˚C | 0 | Fig. 1A suggests no reduction in mean lifespan (no statistics) | Fig. 1B suggests a modest reduction in lifespan (no statistics) | Not reported | NS | (Hars et al., 2007) |
| *daf-2(e1370)* | *lgg-3* (formerly *atg-12*) | *ATG12* | RNAi by feeding | Young adult | 15˚C | 0 | Slight reduction (data not shown) | No effect (data not shown) | Not reported | NS | (Hars et al., 2007) |
| *daf-2(e1370)* | *lgg-3* (formerly *atg-12*) | *ATG12* | RNAi by feeding | Young adult of previous generation (RNAi during development) | 15˚C | 0 | -38.8 ($p$=2.1 x $10^{-13}$) | -35.7 ($p$=1.1 x $10^{-7}$) | Yes, $p$=0.129 (whole populations), but $p$ = 0.0169 (last 50% survivors) | NS | (Hars et al., 2007) |
| *fer-15(b26); daf-2(mu150); fem-1(hc17)* | *bec-1* | *BECN1* | RNAi by feeding | Young adult | 20˚C | 0 | Reduced lifespan in 6 trials ($p<0.05$ in all cases) | No reduction in N2 lifespan in 3 trials (NS) | Yes | 6 | (Hansen et al., 2008) |
| *fer-15(b26); daf-2(mu150); fem-1(hc17)* | *vps-34* | *PIK3C3* | RNAi by feeding | Young adult | 20˚C | 0 | Reduced lifespan in 3 trials ($p<0.05$ in all cases) | No reduction in N2 lifespan in 3 trials (NS) | Yes | 3 | (Hansen et al., 2008) |
| *daf-2(e1370)* | *atg-9* | *ATG9A/ATG9B* | RNAi by feeding | Not specified | 25˚C | 1.21 M FUDR on day 1 of adulthood only | -17.5 (no statistical test) | Not reported | Unknown | NS | (Toth et al., 2008) |
| *daf-2(e1370)* | *atg-18* | *WIPI1/WIPI2* | Mutation, *atg-18(gk378)* | N/A | 25˚C | 1.21 M FUDR on day 1 of adulthood only | -24.6 (no statistical test) | -23.2 ($p<0.001$) | Unlikely (no statistical test) | NS | (Toth et al., 2008) |

| | | | | | | | | | | | |
|---|---|---|---|---|---|---|---|---|---|---|---|
| *daf-2* RNAi | *atg-18* | *WIPI1/WIPI2* | Mutation, *atg-18(gk378)* | L2 larval stage | 25˚C | 1.21 M FUDR on day 1 of adulthood only | -17.8 (no statistical test) | -23.2 ($p$<0.001) | No | NS | (Toth et al., 2008) |
| *daf-2(e1370)* | *bec-1* | *BECN1* | Mutation, *bec-1(ok691); Ex[bec-1(+)]* | N/A | 25˚C | 1.21 M FUDR on day 1 of adulthood only | -18.6 (no statistical test) | -26.4 ($p$<0.002) | No | NS | (Toth et al., 2008) |
| *daf-2(e1370)* | *lgg-1* | *GABARAP* | RNAi by feeding | Not specified | 25˚C | 1.21 M FUDR on day 1 of adulthood only | -8.2 (no statistical test) | Not reported | Unknown | NS | (Toth et al., 2008) |
| *daf-2(e1370)* | *atg-3* | *ATG3* | RNAi by feeding | Maternal RNAi | 20˚C | 800 µM | **+18.3** ($p$=0.012) | -18.6 ($p$=0.0026) | No | 2 | (Hashimoto et al., 2009) |
| *daf-2(e1370)* | *atg-4.1* | *ATG4A/ATG4B* | RNAi by feeding | Maternal RNAi | 20˚C | 800 µM | +0.3 (NS) | -20.1 ($p$<0.0001) | No | 2 | (Hashimoto et al., 2009) |
| *daf-2(e1370)* | *atg-4.2* | *ATG4C/ATG4D* | RNAi by feeding | Maternal RNAi | 20˚C | 800 µM | 0.0 (NS) | -7.1 ($p$=0.035) | No | 2 | (Hashimoto et al., 2009) |
| *daf-2(e1370)* | *atg-5* | *ATG5* | RNAi by feeding | Maternal RNAi | 20˚C | 800 µM | **+26.4** ($p$=0.0014) | -19.2 ($p$=0.0001) | No | 2 | (Hashimoto et al., 2009) |
| *daf-2(e1370)* | *atg-7* | *ATG7* | RNAi by feeding | Maternal RNAi | 20˚C | 800 µM | **+24.9** ($p$=0.0005) | +3.7 (NS) | No | 3 | (Hashimoto et al., 2009) |
| *daf-2(e1370)* | *atg-9* | *ATG9A/ATG9B* | RNAi by feeding | Maternal RNAi | 20˚C | 800 µM | **+21.2** ($p$=0.014) | -7.1 (NS) | No | 2 | (Hashimoto et al., 2009) |
| *daf-2(e1370)* | *atg-10* | *ATG10* | RNAi by feeding | Maternal RNAi | 20˚C | 800 µM | +7.5 (NS) | 3.7 (NS) | No | 2 | (Hashimoto et al., 2009) |
| *daf-2(e1370)* | *atg-16.2* | *ATG16L1/ ATG16L2* | RNAi by feeding | Maternal RNAi | 20˚C | 800 µM | **+17.8** ($p$ = 0.0084) | -7.7 (NS) | No | 2 | (Hashimoto et al., 2009) |
| *daf-2(e1370)* | *atg-18* | *WIPI1/WIPI2* | RNAi by feeding | Maternal RNAi | 20˚C | 800 µM | -35.8 ($p$=0.0011) | -47.8 ($p$<0.0001) | No | 2 | (Hashimoto et al., 2009) |
| *daf-2(e1370)* | *bec-1* | *BECN1* | RNAi by feeding | Maternal RNAi | 20˚C | 800 µM | -23.2 ($p$=0.021) | -16.0 ($p$=0.011) | No | 2 | (Hashimoto et al., 2009) |
| *daf-2(e1370)* | *lgg-1* | *GABARAP* | RNAi by feeding | Maternal RNAi | 20˚C | 800 µM | +1.3 (NS) | -12.3 ($p$=0.0012) | No | 1 | (Hashimoto et al., 2009) |
| *daf-2(e1370)* | *lgg-2* | *LC3* | RNAi by feeding | Maternal RNAi | 20˚C | 800 µM | +10.3 (NS) | 0.0 (NS) | No | 2 | (Hashimoto et al., 2009) |
| *daf-2(e1370)* | *lgg-3* | *ATG12* | RNAi by feeding | Maternal RNAi | 20˚C | 800 µM | **+25.6** ($p$=0.0027) | -2.8 (NS) | No | 2 | (Hashimoto et al., 2009) |
| *daf-2(e1370)* | *unc-51* | *ULK1/UKL2* | RNAi by feeding | Maternal RNAi | 20˚C | 800 µM | **+14.8** ($p$=0.032) | +0.2 (NS) | No | 3 | (Hashimoto et al., 2009) |
| *daf-2(e1370)* | *atg-3* | *ATG3* | RNAi by feeding | Early adulthood | 20˚C | 800 µM | +1.1 (NS) | -0.2 (NS) | No | 2 | (Hashimoto et al., 2009) |
| *daf-2(e1370)* | *atg-4.1* | *ATG4A/ATG4B* | RNAi by feeding | Early adulthood | 20˚C | 800 µM | -8.6 ($p$=0.022) | -3.9 (NS) | Possible (no statistical test) | 2 | (Hashimoto et al., 2009) |

| | | | | | | | | | | | |
|---|---|---|---|---|---|---|---|---|---|---|---|
| *daf-2(e1370)* | *atg-4.2* | *ATG4C/ATG4D* | RNAi by feeding | Early adulthood | 20˚C | 800 µM | -2.2 (NS) | -5.8 (NS) | No | 2 | (Hashimoto et al., 2009) |
| *daf-2(e1370)* | *atg-5* | *ATG5* | RNAi by feeding | Early adulthood | 20˚C | 800 µM | -0.7 (NS) | +2.2 (NS) | No | 2 | (Hashimoto et al., 2009) |
| *daf-2(e1370)* | *atg-7* | *ATG7* | RNAi by feeding | Early adulthood | 20˚C | 800 µM | -2.55 (NS) | **+7.6** (*p*=0.020) | No | 3 | (Hashimoto et al., 2009) |
| *daf-2(e1370)* | *atg-9* | *ATG9A/ATG9B* | RNAi by feeding | Early adulthood | 20˚C | 800 µM | **+15.8** (*p*=0.0013) | **+11.4** (*p*=0.0013) | No | 3 | (Hashimoto et al., 2009) |
| *daf-2(e1370)* | *atg-10* | *ATG10* | RNAi by feeding | Early adulthood | 20˚C | 800 µM | -3.0 (NS) | -2.0 (NS) | No | 2 | (Hashimoto et al., 2009) |
| *daf-2(e1370)* | *atg-16.2* | *ATG16L1/ ATG16L2* | RNAi by feeding | Early adulthood | 20˚C | 800 µM | +1.4 (NS) | +3.4 (NS) | No | 2 | (Hashimoto et al., 2009) |
| *daf-2(e1370)* | *atg-18* | *WIPI1/WIPI2* | RNAi by feeding | Early adulthood | 20˚C | 800 µM | -4.3 (NS) | -22.9 (*p*=0.0010) | No | 2 | (Hashimoto et al., 2009) |
| *daf-2(e1370)* | *lgg-1* | *GABARAP* | RNAi by feeding | Early adulthood | 20˚C | 800 µM | -3.3 (NS) | -10.4 (*p*=0.019) | No | 2 | (Hashimoto et al., 2009) |
| *daf-2(e1370)* | *lgg-2* | *LC3* | RNAi by feeding | Early adulthood | 20˚C | 800 µM | +6.2 (NS) | +5.3 (NS) | No | 2 | (Hashimoto et al., 2009) |
| *daf-2(e1370)* | *lgg-3* | *ATG12* | RNAi by feeding | Early adulthood | 20˚C | 800 µM | -1.4 (NS) | +8.1 (NS) | No | 2 | (Hashimoto et al., 2009) |
| *daf-2(e1370)* | *bec-1* | *BECN1* | RNAi by feeding | Early adulthood | 20˚C | 800 µM | **+11.4** (*p*=0.0029) | **+10.1** (*p*=0.028) | No | 3 | (Hashimoto et al., 2009) |
| *daf-2(e1370)* | *unc-51* | *ULK1/UKL2* | RNAi by feeding | Early adulthood | 20˚C | 800 µM | **+11.0** (*p*=0.010) | **+9.4** (*p*=0.024) | No | 3 | (Hashimoto et al., 2009) |
| *daf-2(e1370)* | *atg-18* | *WIPI1/WIPI2* | RNAi by feeding | Early adulthood | 20˚C | 0 | -56, -51, -51, -52, -52 (*p*<0.0001) | Test not performed | Insufficient data | 5 | (Chang et al., 2017) |
| *daf-2(e1370* | *lgg-1* | *GABARAP* | RNAi by feeding | Early adulthood | 20˚C | 0 | -21, -22 (*p*<0.0001) | Test not performed | Insufficient data | 2 | (Chang et al., 2017) |
| *daf-2(e1370)* | *atg-18* | *WIPI1/WIPI2* | Mutation, *atg-18(gk378)* | N/A | 20˚C | 0 | -39, -25, -29 (median lifespan) (*p*<0.0001) | -50, -33, -32 (median lifespan) (*p*<0.0001) | Possible trend (no statistical test) | 3 | (Minnerly et al., 2017) |
| *rrf-3(pk1426); daf-2(e1370)* | *bec-1* | *BECN1* | RNAi by feeding | Day 10 of adulthood[3] | 20˚C | 0 | **+21.3, +11.3, +17.7** (*p*<0.0001) | **+39.3, +51.9, +62.5** (*p*<0.0001) | No | 3 | (Wilhelm et al., 2017) |
| *daf-2(e1370)* | *sqst-1* | *SQSTM1* | Mutation, *sqst-1(ok2892)* | N/A | 20˚C | 0 | -10.4 (no statistical test) | +2.0 (*p*=0.6) | Not reported | 1 | (Kumsta et al., 2019) |

[1]Bold, significant increase in lifespan after autophagy pathway gene RNAi.

[2]Number of trials. NS, not specified.

[3]Day 0 defined as L4 stage.

### Supplementary Table 8: Previous reports of effects of inhibition of autophagy on GSC(-) Age

| ***C. elegans* genotype** | ***atg* gene** | **Mammalian gene** | **Form of gene inhibition** | **Age at initiation of RNAi** | **Temperature** | **FUDR concentration (mM)** | **Change in GSC(-) mean lifespan relative to RNAi control (p, log rank)** | **Change in N2 control mean lifespan relative to RNAi control (p, log rank)** | **Greater reduction in lifespan in GSC(-)?** | ***N*[2]** | **Source** |
|---|---|---|---|---|---|---|---|---|---|---|---|
| *glp-1(e2141)* | *bec-1* | *BECN1* | RNAi by feeding | D1 of adulthood | 25˚C until D1, then 20˚C | 0 | -22%, -17%, -21% (*p*<0.0001, 0.0005, <0.0001) | +7%, 0,% +7% (*p*=0.14, 0.94, 0.27) | Yes | 3 | (Lapierre et al., 2011) |
| *glp-1(e2141)* | *lgg-1* | *GABARAP* | RNAi by feeding | D1 of adulthood | 25˚C until D1, then 20˚C | 0 | -18%, -18%, -39% (*p*=0.009, <0.0001, <0.0001) | +1%, +9%, -2% (*p*=0.89, 0.06, 0.30) | Yes | 3 | (Lapierre et al., 2011) |
| *glp-1(e2141)* | *unc-51* | *ULK1/UKL2* | RNAi by feeding | D1 of adulthood | 25˚C until D1, then 20˚C | 0 | -27%, -11%, -17% (*p* <0.0001, 0.040, 0.0028) | +2%, +10%, -2% (*p*=0.90, 0.068, 0.67) | Yes | 3 | (Lapierre et al., 2011) |
| *glp-1(e2141)* | *vps-34* | *PIK3C3* | RNAi by feeding | D1 of adulthood | 25˚C until D1, then 20˚C | 0 | -25%, -26%, -24% (*p*<0.0001) | +6%, -2%, -5% (*p*=0.19, 0.27, 0.52) | Yes | 3 | (Lapierre et al., 2011) |
| *glp-1(e2141)* | *atg-18* | *WIPI1/WIPI2* | RNAi by feeding | D1 of adulthood | 25˚C until D1, then 20˚C | 0 | -35%, -33%, -28% (*p*<0.0001) | +6%, -3%, +10% (*p*=0.38, 0.52, 0.85) | Yes | 3 | (Lapierre et al., 2011) |
| *mes-1(bn17)* | *vps-34* | *PIK3C3* | RNAi by feeding | D1 of adulthood | 25˚C until D1, then 20˚C | 0 | -31% (*p*<0.0001) | +2% (*p*=0.96) | Yes | 3 | (Lapierre et al., 2011) |
| *mes-1(bn17)* | *atg-18* | *WIPI1/WIPI2* | RNAi by feeding | D1 of adulthood | 25˚C until D1, then 20˚C | 0 | -30% (*p*<0.0001) | +3% (*p*=0.60) | Yes | 3 | (Lapierre et al., 2011) |
| *glp-1(e2141)* | *lmp-1* | *LAMP-1* | RNAi by feeding | D1 of adulthood | 25˚C until D1, then 20˚C | 0 | -11%, -4%, -11% (*p*=0.069, 0.58, 0.041) | +1%, -12%, +13% (*p*=0.79, 0.0059, 0.021) | Possibly | 3 | (Lapierre et al., 2013) |
| *glp-1(e2141)* | *vha-16* | *ATP6V0D1* | RNAi by feeding | D1 of adulthood | 25˚C until D1, then 20˚C | 0 | -47%, -42%, -32% (*p*<0.0001) | -11%, -16%, -9% (*p* = 0.0015, 0.0001, 0.011) | Yes | 3 | (Lapierre et al., 2013) |
| *glp-1(e2141)* | *bec-1* | *BECN1* | RNAi by feeding | D10 of adulthood[2] | 25˚C until D1[3], then 20˚C | 0 | **+24.0**, +**34.9**, +**32.5**, +**39.0** (*p*<0.0001) | Not performed | No | 4 | (Wilhelm et al., 2017) |
| *glp-1(e2144)* | *sqst-1* | *SQSTM1* | Mutation, *sqst-1(ok2892)* | N/A | 25˚C until D1, then 20˚C | 0 | -9.9%, -5.5%, -3.3% (no statistical test) | +11%, -11%, -8% (*p*=0.7, 0.003, 0.3) | Not reported | 3 | (Kumsta et al., 2019) |

[1]Bold, significant increase in lifespan after autophagy pathway gene RNAi.
[2]Number of trials. NS, not specified.
[3]Day 0 defined as L4 stage.
